# Recognition of HIV-1 Capsid Licenses Innate Immune Response to Viral Infection

**DOI:** 10.1101/2022.01.10.472699

**Authors:** Sunnie M Yoh, João I. Mamede, Derrick Lau, Narae Ahn, Maria T Sánchez-Aparicio, Joshua Temple, Andrew Tuckwell, Nina V. Fuchs, Gianguido C. Cianci, Laura Riva, Heather Curry, Xin Yin, Stéphanie Gambut, Lacy M. Simons, Judd F. Hultquist, Renate König, Yong Xiong, Adolfo García-Sastre, Till Böcking, Thomas J. Hope, Sumit K. Chanda

**Affiliations:** Sanford Burnham Prebys Medical Discovery Institute, 10901 N. Torrey Pines Rd., La Jolla, CA 92116; Department of Cell and Developmental Biology, Feinberg School of Medicine, Northwestern University, 303 E superior., Chicago, IL60611; EMBL Australia Node in Single Molecule Science, School of Medical Sciences, University of New South Wales, 2052 Kensington, Sydney, Australia; Department of Microbiology, Icahn School of Medicine at Mount Sinai, 1 Gustave L. Levy Place, New York, NY10029; Yale University, 333 Cedar Street, New Haven, CT 06520-8024; Host Pathogen Interaction (NG3), Paul-Ehrlich-Institut, Paul-Ehrlich-Strasse 51-59 Langen, Germany, 63225; Department of Microbial Pathogens and Immunity, Rush University Medical Center, Chicago, IL, USA; Global Health and Emerging Pathogens Institute, Icahn School of Medicine at Mount Sinai, 1 Gustave L. Levy Place, New York, NY10029; Department of Medicine, Division of Infectious Diseases, Icahn School of Medicine at Mount Sinai, 1 Gustave L. Levy Place, New York, NY10029; Calibr, a division of The Scripps Research Institute, 11119 North Torrey Pines Road, La Jolla, CA92037; State Key Laboratory of Veterinary Biotechnology, Harbin Veterinary Research Institute, Chinese Academy of Agricultural Sciences, Harbin 150069, P.R. China; Department of Medicine, Division of Infectious Diseases, Northwestern University Feinberg School of Medicine, Chicago, IL, 60611, USA; Center for Pathogen Genomics and Microbial Evolution, Institute for Global Health, Northwestern University Feinberg School of Medicine, Chicago, IL 60611, USA

**Author notes:** Correspondence &. Equal contribution (Sunnie M Yoh, João I. Mamede). Equal contribution (Derrick Lau, Narae Ahn).

**Keywords:** PQBP1, cGAS, HIV-1 Capsid, Innate sensing, Two-factor authentication, Uncoating

## Abstract

Cyclic GMP-AMP synthase (cGAS) is a primary sensor of aberrant DNA that governs an innate immune signaling cascade, leading to the induction of the type-I interferon response. We have previously identified polyglutamine binding protein 1, PQBP1, as an adaptor molecule required for cGAS-mediated innate immune response of lentiviruses, including the human immunodeficiency virus 1 (HIV-1), but dispensable for the recognition of DNA viruses. HIV-1- encoded DNA is synthesized as a single copy from its RNA genome, and is subsequently integrated into the host chromatin. HIV-1 then produces progeny through amplification and packaging of its RNA genome, thus, in contrast to DNA viruses, HIV-1 DNA is both transient and of low abundance. However, the molecular basis for the detection and verification of this low abundance HIV-1 DNA pathogen-associated molecular pattern (PAMP) is not understood. Here, we elucidate a two-factor authentication strategy that is employed by the innate immune surveillance machinery to selectively respond to the low concentration of PAMP, while discerning these species from extranuclear DNA molecules. We find that, upon HIV-1 infection, PQBP1 decorates intact viral capsid, which serves as a primary verification step for the viral nucleic acid cargo. As the reverse transcription and capsid disassembly initiate, cGAS protein is then recruited to the capsid in a PQBP1-dependent manner, enabling cGAS molecules to be co-positioned at the site of PAMP generation. Thus, these data indicate that PQBP1 recognition of the HIV-1 capsid sanctions a robust cGAS-dependent response to a limited abundance and short-lived DNA PAMP. Critically, this illuminates a molecular strategy wherein the modular recruitment of co-factors to germline encoded pattern recognition receptors (PRRs) serves to enhance repertoire of pathogens that can be sensed by the innate immune surveillance machinery.

## INTRODUCTION

The cGAS signaling pathway has been established as a critical regulator of the innate immune response to cytoplasmic DNA (Ablasser and Chen, 2019; Chin, 2019; Reinert et al., 2016). Signaling can be induced by cytosolic delivery of exogenous double stranded (ds) DNA longer than 50 nucleotides in length (Gao et al., 2013; Paludan and Bowie, 2013). Upon binding to dsDNA, cGAS synthesizes cyclic GMP-AMP dinucleotide (cGAMP) which serves as a second messenger to induce STING-mediated IRF3/type-I IFN signaling (Gao et al., 2013). Interestingly, cGAS can also sense lentiviral infections, but unlike DNA viruses, requires an adaptor protein, PQBP1 (Yoh et al., 2015). Recently, the non-POU domain-containing octamer-binding protein (NONO) has been implicated in the innate sensing of nuclear HIV DNA (Lahaye et al., 2018).

The HIV-1 capsid, a protein shell composed of oligomeric structural capsid proteins (CA), encapsulates viral proteins and the RNA genome (Rankovic et al., 2017). After the virion entry into cytosol, capsid initiates the progressive disassembly of CA through a poorly defined process of uncoating (Campbell and Hope, 2015; Pornillos et al., 2011). The CA structure continues to disassemble as the viral reverse transcription complex (RTC) migrates to the nucleus (Burdick et al., 2020; Hulme et al., 2011; Mamede et al., 2017; Sood et al., 2017). The initiation of capsid disassembly process is linked to the early steps of reverse transcription, which commences with the first (-) strand cDNA synthesis and is completed with second (+) strand synthesis (Christensen et al., 2020; Hu and Hughes, 2012; Hulme et al., 2011; Mallery et al., 2018; Mamede et al., 2017; Rankovic et al., 2018; Rankovic et al., 2017; Soliman et al., 2017). The HIV-1 DNAs can serve as a pathogen-associated molecular pattern (PAMP) to activate cGAS (Cosnefroy et al., 2016; Doitsh et al., 2010; Yoh et al., 2015). Upon the completion of reverse transcription, the DNA is then integrated into the host genome, where it becomes indistinguishable from chromosomal DNA. Only viral RNA nucleic acid species, biochemically indistinct from host RNAs, are present during subsequent steps of the viral life cycle. Thus, in contrast to DNA viruses which amplify copies of their DNA genomes upon infection, HIV-1 only produces one copy of DNA molecule per reverse transcription competent virion (Hu and Hughes, 2012), and its availability for innate sensing is temporally and spatially constricted as the HIV RTC transits to the nucleus and HIV DNA integrates into host chromatin. The molecular basis of how immune system can sense these transient and low-copy lentiviral DNA species, but not low-abundance self-DNAs, including extranuclear DNAs from mitochondrial or nuclear leakage, is not completely understood.

In this report, we find that the innate sensing machinery employs a unique molecular strategy to license innate immune response through a two-step authentication process. First, retrovirus-specific innate co-sensor PQBP1 specifically recognizes intact capsids of incoming HIV-1 viral particles. Subsequently, disassembly of the viral capsid triggers the PQBP1- dependent recruitment of cGAS in a NONO-independent manner, enabling enzymatic activation of the sensor upon the initiation of HIV DNA production. Thus, PQBP1 recognition of the HIV-1 capsid is the proximal event that serves to distinguish its cargo from self-nucleic acids, through licensing the recruitment of cGAS at the site of PAMP generation. Importantly, this molecular strategy reveals that modular engagement of co-factors to PRRs can enable the innate immune surveillance machinery to respond to an enhanced repertoire of pathogen-encoded PAMPs, while limiting deleterious responses to host genome-encoded DNA molecules.

## RESULTS

### PQBP1 colocalizes with capsid of incoming virions during the early steps of infection

To better understand the spatial dynamics of HIV-1 innate immune recognition and response, we utilized super-resolution 3D-SIM microscopy to assess PQBP1 association with incoming HIV-1 virions. Briefly, PMA-differentiated THP-1 cells (PMA-THP-1) were infected with HIV-1 viruses labeled with Gag-Integrase (IN)-mRuby3 (Dharan et al., 2017; Hulme et al., 2015; Mamede et al., 2017) and one hour post fusion, the cells were fixed and stained by immunofluorescence (IF) against PQBP1 protein (green; Figure 1A). Colocalization of PQBP1 with virus particles (IN- mRuby3; red) was assessed by measuring the distance from each IN-labeled virus puncta to its nearest neighbor PQBP1 dot centroid (Figure 1B; Figure S1). The distribution of IN-PQBP1 nearest neighbor distances (*d*) is shown in blue (Figure 1B), which harbored a sharp peak at shorter distances (∼0.12 µm) and a broad shoulder at distances larger than 0.5 µm, indicating two distinctive populations. For comparison, we performed *in silico* randomized labeling of IN and PQBP1 signals and calculated the *d* distribution (magenta line, left, Figure 1B). The randomized distribution displayed a single broad peak at 0.8-0.9 µm, which indicates the g PQBP1-IN colocalization peak is unlikely due to stochasticity, while the observed broad shoulder (0.5 µm) most likely represents INs that are not colocalizing with PQBP1. To quantify the observed colocalization, we compared the cumulative probability distribution of *d* across a range of thresholds to ascertain colocalization frequencies (right, Figure 1B; Figure S1D; see Supplemental Text for detail). At a threshold of 0.4 µm, we find that more than 40% of total INs are colocalizing with PQBP1, compared to less than 9% for the randomized data (right, Figure 1B). Taken together, at thresholds established below 1 µm, these data indicate that PQBP1 significantly colocalizes with incoming HIV-1 virions.

**Figure 1.**
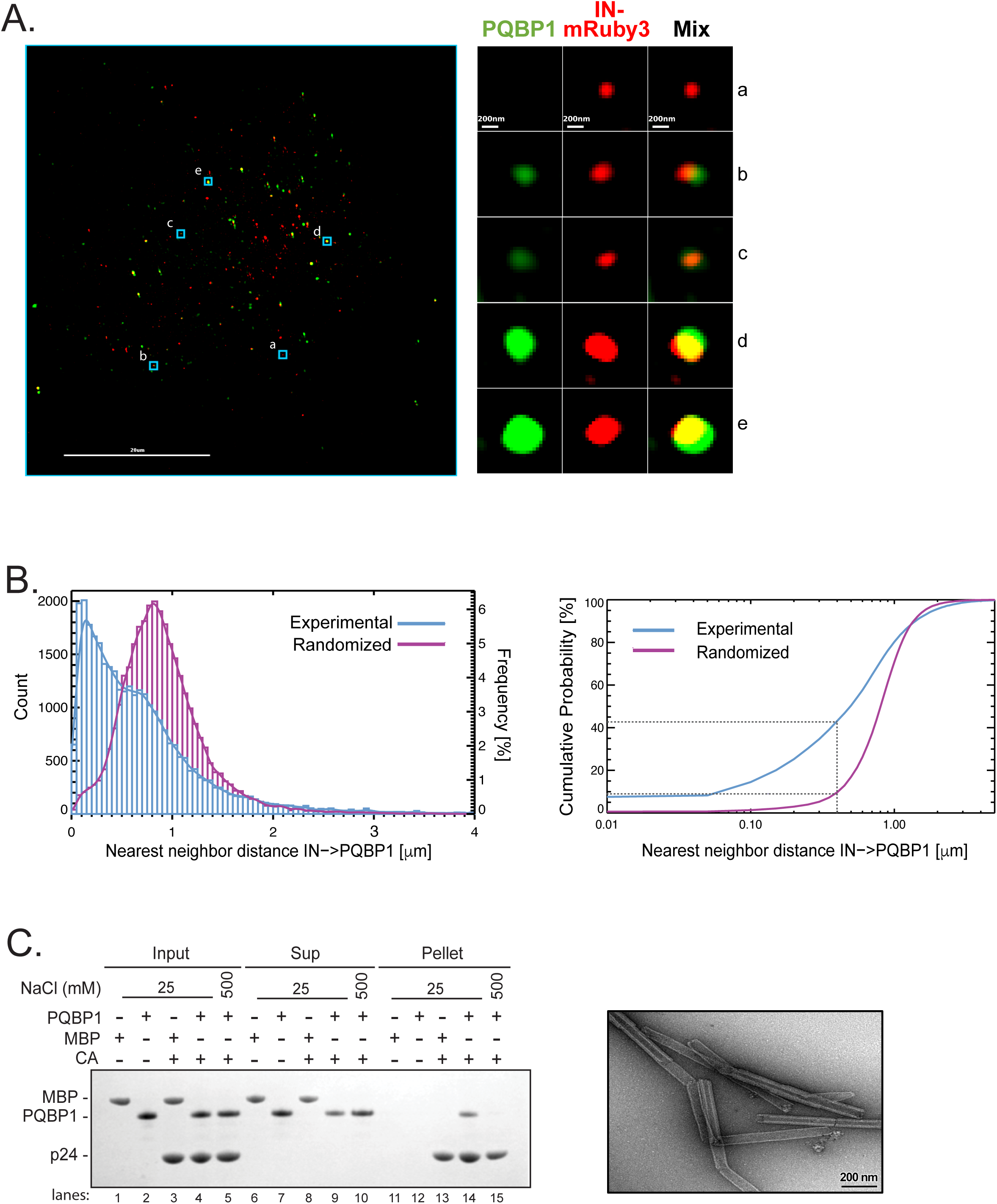
PQBP1 colocalizes with capsid of incoming virions during the early steps of infection. (A) Single Z image of PMA-differentiated THP-1 cells (PMA-THP-1) infected with HIV- 1 virions labeled with Gag-IN-mRuby3 (red). The viral fusion was synchronized and one-hour post-infection, immunostaining of PQBP1 protein (green) was performed. Zoomed images of individual viral particles associating with PQBP1 are shown on the right. (B) Left, distribution of IN-to-PQBP1 nearest neighbor distances (Count). The distribution of experimental data (blue histogram) is compared with the one generated from *in silico* randomized dots (magenta histogram). A kernel density estimate of each distribution is overplotted as a solid curve, represented as Frequency, N=32,033 INs. Right, the cumulative probability of nearest neighbor distance (*d*) measures the percentage of IN dots that have PQBP1 within a defined *d*. For example, the percentage of INs that have PQBP1 at *d* < 0.4 µm is a 43% of for experimental data and 9% for randomized dots as highlighted by grey dotted lines. (C) CA tube co-pelleting assay. Insoluble cross-linked CA A14C/E45C tubes were incubated together with PQBP1 or maltose binding protein (MBP; Input) and separated into supernatant and pellet fractions (Sup and Pellet, respectively) then analyzed via reducing SDS-PAGE. The data are representative of at least three independent experiments.

HIV-1 capsid is composed of ∼1500 copies of CA monomers that self-assemble into lattices of hexamers and pentamers to form fullerene-like cones (Perilla and Gronenborn, 2016; Pornillos et al., 2011; Summers et al., 2019). The lattice serves as a binding platform for numerous cellular factors; thus, HIV-1 capsid defines a critical host-pathogen interface after cellular entry (Campbell and Hope, 2015; James and Jacques, 2018; Novikova et al., 2019; Summers et al., 2019). Since PQBP1 was found to colocalize with incoming virus particles, we examined if there was a direct interaction between recombinant purified PQBP1 and cross-linked CA tubes that recapitulates the hexameric lattice found in virions [right, Figure 1C; (Mattei et al., 2016)]. Due to their size (∼50 nm in diameter and ∼500 nm in length), CA tubes become insoluble upon assembly, which can be utilized to probe a cellular factor binding by co-sedimentation [left, Figure 1C; (Summers et al., 2019)]. We find that PQBP1, but not a negative control maltose binding protein (MBP), specifically co-pelleted with CA tubes (compare lane 13 & 14), and this association could be disrupted in high-salt condition (compare lanes 14 & 15). These data indicate a biochemical association of PQBP1 for CA tubes that corroborates their colocalization within the infected cells.

### PQBP1 directly interacts with HIV-1 capsids through its amino-terminus

To date, at least four binding sites on the capsid have been identified for host factor binding: (1) the CypA binding loop (residues 85-95, also recognized by Nup358), (2) the FG-binding site located between adjacent CA subunits centered at residue 74 in the hexamer recognized by CPSF6 and Nup153, (3) the R18 residues of six CA subunits that forms an electropositive pore in the center of the hexamer/pentamer as a binding site for polyanions including dNTPs, IP6 and the host protein FEZ1, and (4) the electronegative inter-hexamer junction at the three fold symmetry of the capsid lattice that recruits the antiviral protein MxB (Bhattacharya et al., 2014; Gamble et al., 1996; Mallery et al., 2018; Price et al., 2014; Smaga et al., 2019). We then sought to delineate the binding interface between capsid and PQBP1 utilizing two-color coincidence detection (TCCD). This technique measures fluorescence intensity fluctuations from both cross- linked CA A204C self-assembled particles, labeled with AF568, and capsid-binding proteins, labeled with AF488, diffusing through the confocal volume (Figure 2A, orange and green traces respectively; Figure S2;(Lau et al., 2021; Lau et al., 2019)). Accumulation of binders on the capsid results in the appearance of fluorescence peaks in the binder trace that coincide with the capsid peak (Figure 2A, middle; Figure S2B), whereby the variation in the peak amplitudes is due to the heterogeneity in the size of the *in vitro* assembled capsid particles. As a control, we assessed the binding of a CPSF6 peptide comprising the capsid-binding residues (Bhattacharya et al., 2014; Lau et al., 2019; Price et al., 2014) to CA A204C particles (Figure 2B). Consistent with previously published data, we observed a significant reduction in CPSF6 binding to the N74D capsid mutant that disables the FG binding site in comparison to the strong binding to WT and R18G capsid mutant (Mallery et al., 2018). Additionally, the CPSF6-capsid interaction was not competed off by hexacarboxybenzene (HCB), a polyanion that binds to the R18 ring (HCB; Figure 2A, bottom, Figure 2B, Figure S2C; (Jacques et al., 2016)). In contrast, fluorescein-dATP, which also binds to R18 ring, was competed off in the presence of HCB and failed to associate with R18G mutant capsid (Figure 2B, Figure S2; (Jacques et al., 2016)). Importantly, we observed that PQBP1, which was directly conjugated with AF488 at the C60 residue, bound robustly to labeled CA A204C particle, as well as the CA A204C particles bearing the N74D mutation; however, PQBP1 was effectively competed off by HCB and failed to bind to the R18G mutant (Figure 2B, Figure S2C). Lastly, we find that the N-terminal domain (residues 1 to 46) of PQBP1, which has a cluster of negatively charged residues, showed greater accumulation on CA particles than the C-terminal portions of the protein (residues 47-265; Figure 2C; Figure S2C). Taken together, these results suggest that the N-terminal region of PQBP1 and the ring of R18 residues of the capsid make critical contributions to the viral-host interface.

**Figure 2.**
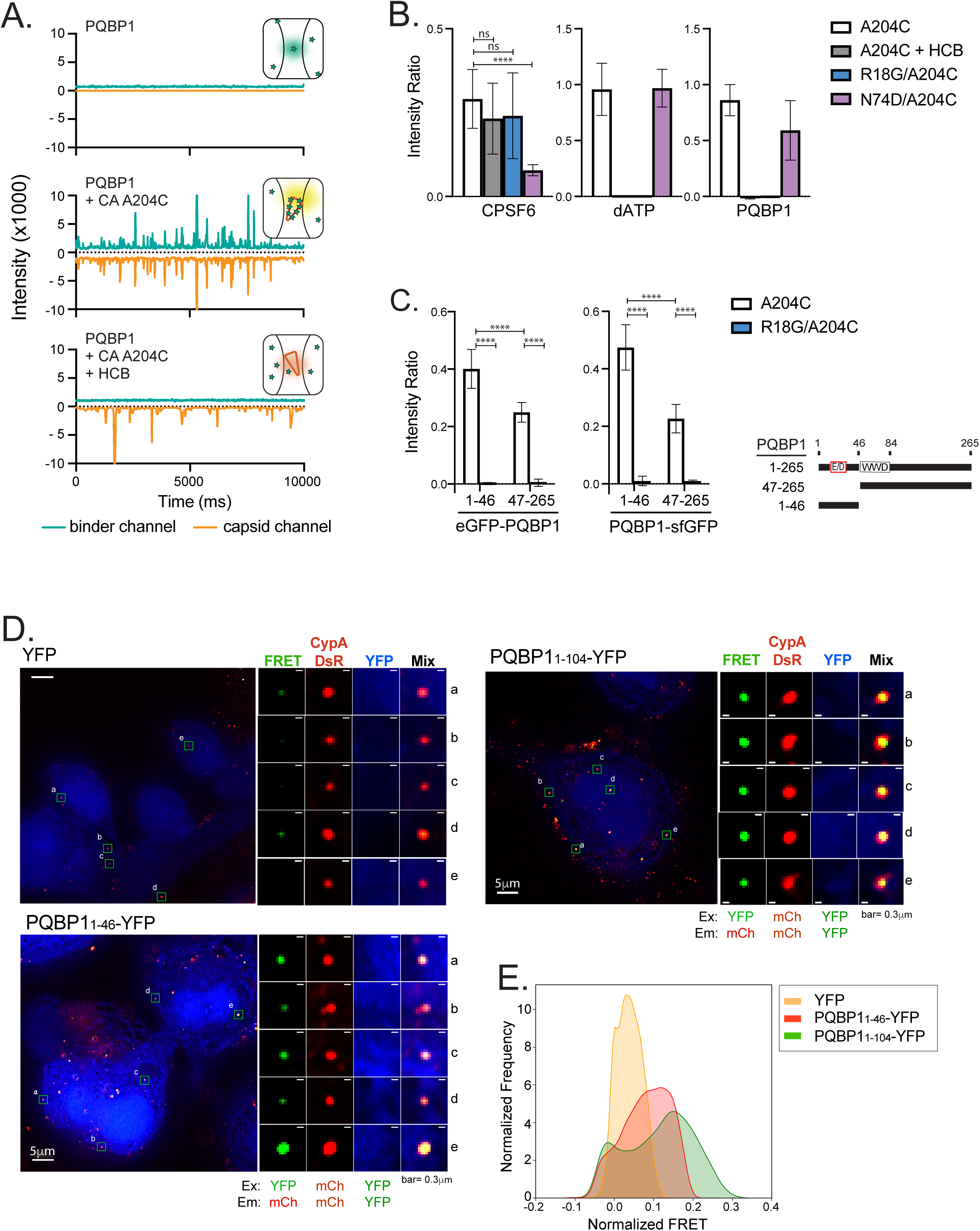
PQBP1 directly interacts with HIV-1 capsids through its amino-terminus. (A) Representative TCCD traces of AF488-PQBP1 (green) and AF568-labeled CA A204C particles (orange). The insets show schematics of the species detected by TCCD with PQBP1 and CA A204C particles represented as green stars and orange cones, respectively. Top, featureless PQBP1 trace due to the diffusion of PQBP1 monomers; middle, coincident peaks in both traces due to co-diffusion of multiple PQBP1 molecules bound to CA particles; bottom, featureless PQBP1 trace due to non-associated diffusion of PQBP1 monomers and CA particles in the presence of HCB. (B) PQBP1 binds to the R18 pore of capsid. TCCD analysis of AF488-PQBP1, AF488-CPSF6_313-327_ and fluorescein-dATP to CA A204C particles in the absence and presence of HCB, and to R18G or N74D CA particles. (C) The N-terminal 46 residues of PQBP1 bound strongly to capsid. TCCD binding analysis of GFP fusions of PQBP1_1-46_ and PQBP1_47-265_ to CA A204C particles in the absence and presence of HCB. Data are representative of at least two independent experiments. One-way ANOVA, **** p<0.0001, ns=no significance. (D) Fluorescence resonance energy transfer (FRET) assay to visualize interaction between PQBP1 and capsid of incoming virions. PMA-THP-1 cells stably expressing either eYFP, PQBP1__1-46_- eYFP or PQBP1_1_-104_-eYFP (blue) were infected with HIV-1 packaged with CypA-DsRed (red) for 1.5 hours, followed by PFA fixation, imaging, and FRET analysis. Representative images and distributions of FRET values normalized against an uninfected counterpart were shown (see Method for detail). FRET excitation and emission wavelengths for YFP and mCherry are as annotated. R0 calculated to be 60.98Å (https://www.fpbase.org/fret/). Data are representative of at least two independent experiments.

To corroborate these *in vitro* binding studies, the association between capsid binding domain of PQBP1 and incoming viral cores in the infected cells was assessed by a fluorescence resonance energy transfer (FRET) assay. The strong affinity of CypA-DsRed protein to the HIV- 1 capsid enables the protein to be specifically packaged into virions during the viral production and remains bound to the capsid surface post fusion (Francis et al., 2016; Francis and Melikyan, 2018; Sood et al., 2017). We reasoned that if the PQBP1-eYFP fusion protein is indeed recruited to the capsid, it should be in close proximity (< 10 nm) to capsid bound CypA-DsRed, allowing for FRET between the eYFP and DsRed fluorophores. Indeed, THP-1 cells stably expressing eYFP fused PQBP 1-104 or 1-46 proteins (blue) showed higher FRET signals (green) compared to YFP alone (Figure 2D). The FRET signal of each infected sample was normalized by the background signal from the corresponding uninfected sample [(Francis et al., 2016; Francis and Melikyan, 2018; Sood et al., 2017); see Star Method]. The distribution of normalized FRET values of both PQBP1 expressing cells were clearly distinctive and shifted to higher values from that of cells expressing eYFP alone (Figure 2E). As a control, we observed that the distribution of DsRed signals was equivalent among the infected THP-1 cells being subjected to the analysis confirming comparable infection level (Figure S2D). This observation recapitulates the in vitro association of PQBP1and cGAS in the cytoplasm during the infection. In summary, these data from orthogonal assays support that PQBP1 directly associates with the assembled capsid and this interaction is likely mediated through the N-terminus of PQBP1.

### PQBP1 is required for cGAS recruitment to incoming virus particles

We have previously demonstrated that a physical interaction of PQBP1 with cGAS is essential for the innate sensing of HIV-1 infection (Yoh et al., 2015). We hypothesized that cGAS is tethered to incoming virus particles via PQBP1 directly associating with the viral capsid. To address this hypothesis, we have performed super-resolution 3D-SIM IF to exam cGAS association with incoming virus particles during the early-state of post-fusion viral infection (Figure 3A; Figure S3A; see Method). Assessing the puncta overlap between IN-mRuby3 (virus) and the associating proteins, we observed only a fraction of INs co-associated with cGAS upon 1 hr of post-infection (∼8% on average compared to ∼60 % for PQBP1); yet, majority of cGAS molecules associating with INs were also associated with PQBP1 (bottom, Figure 3A). Next, we examined the kinetics of PQBP1 and cGAS recruitment to incoming virus particles (RTCs), utilizing a virus containing GFP as the fluid phase marker (HIV-iGFP) to monitor the core after the fusion. Soon after the fusion of the virus with the cellular membrane, the HIV-1 core changes in a way that allows the encapsulated GFP to leak out due to the loss of core integrity (Chen et al., 2007; Hubner et al., 2007; Hubner et al., 2009; Mamede et al., 2017). This loss of integrity is dependent on the process of reverse transcription where the blocking of the initiation or the first strand transfer of the reverse transcription delays the initiation of capsid disassembly (Cosnefroy et al., 2016; Mamede et al., 2017). We utilized this system to stage the progression of the cytoplasmic HIV complexes and the state of capsid. Upon the loss of capsid integrity, the viral DNAs being generated by ongoing reverse transcription are likely to become accessible to cGAS. By infecting MDDCs with the HIV- 1 virus dual-labeled with iGFP and Gag-Integrase (IN)-mRuby3, we combined live cell imaging to monitor the state of the capsid core of each virus particle within the infected cells, followed by timed fixation and immuno-staining against PQBP1 and cGAS to determine the level of their association with each RTC (Figure 3B). For the analysis, each intracellular (IN)-mRuby3 complex was categorized based on the characteristic iGFP stepwise signal decay: (1) unfused (no loss of iGFP), (2) fused but retaining an intact capsid (partial loss of iGFP), and (3) after fusion with loss of capsid integrity (complete loss of iGFP) (Video S1; Figure S3B; iGFP panel, Figure 3B). PQBP1 and cGAS association with each virus particle was then quantified by post-fixation immunostaining of PQBP1 and cGAS, followed with spatial correlation to IN foci (individual panels and quantification graphs in Figure 3B & Figure S3B). We found that PQBP1 associated with post-fusion particles regardless of whether the capsid was intact or open, while most cGAS co- localized only after loss of capsid integrity.

**Figure 3.**
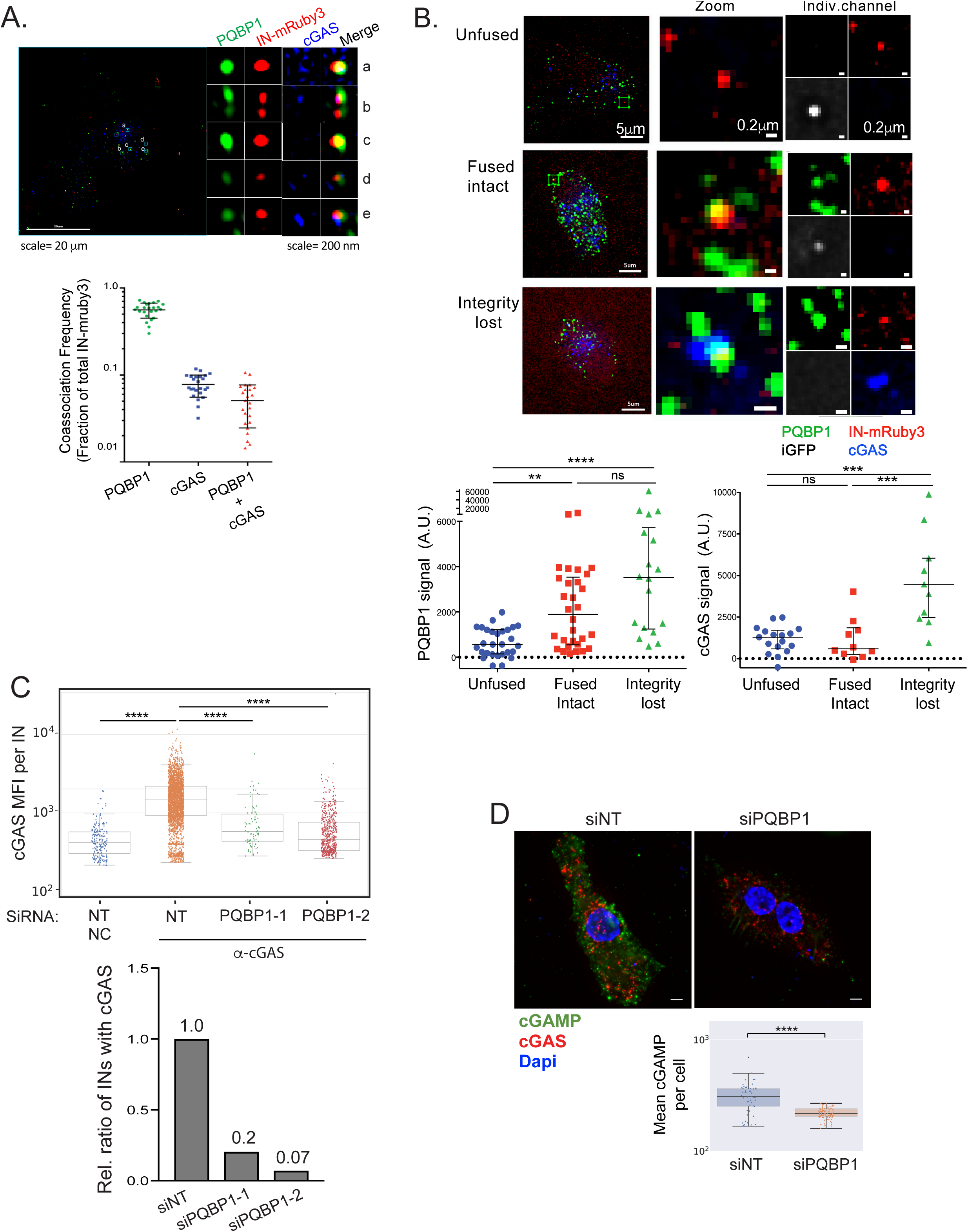
PQBP1 is required for cGAS recruitment to incoming virus particles. (A) cGAS and PQBP1 association with incoming viral particles. Single Z slice images of PMA-THP-1 cells infected with HIV-1 labeled with Gag-IN-mRuby3 (red) in the presence of VLP-Vpx coinfection for 1 hr, followed by antibody detection of PQBP1 protein (green) and cGAS (blue). Detailed cropped images of individual viral complexes associating with PQBP1 and cGAS at different levels and distributions are shown on the right. Fractions of total IN-mRuby3 foci showing association with PQBP1, cGAS (independently) and simultaneously positive for signals of viral particle, PQBP1, and cGAS are quantified at the bottom. (B) PQBP1 recruitment to viral particles precedes cGAS recruitment. MDDCs were infected with HIV-1 labeled with iGFP fluid phage marker and IN- mRuby3 at low MOI and subjected to a time-lapse imaging for an hour, followed by fixed IF for PQBP1 and cGAS. Signal intensity of PQBP1/cGAS for each viral particle are quantified and plotted according to the structural state of capsids, assessed by iGFP signal status (see Method, bottom graphs). Kruskal Wallis H; KW p<0.001), followed by Dunn’s multiple comparisons *** p<0.001, ** p<0.01. Data represent sum of 2 to 3 independent experiments. (C) PQBP1 is required for cGAS recruitment to viral particles. Top, PMA-THP-1, treated with either a non-targeting siRNA (NT) or siRNAs targeting PQBP1 (PQBP1-1 and -2), were infected with HIV-1 viruses (Gag-IN- mRuby3) for 2.5 hours, followed by fixed imaging for cGAS. NC denotes negative control where the cells were only stained with secondary antibodies. Mean fluorescence intensity (MFI) of cGAS signal per viral particle (IN) is quantified. Median and error bar (-/+ 1.5*IQR) are shown. Box indicates the interquartile range (IQR). Dunn’s column comparison **** p<0.0001. Bottom, relative ratio of viral particles (INs) where co-associating cGAS MFI signals above the maximum value of NC sample are plotted from the data shown in top graph. All the HIV-1 infections of PMA-THP-1 cells were performed in the presence of VLP-Vpx co-infection. (D) Knockdown of PQBP1 results in decreased cGAMP production. PMA-THP-1, transfected with non-targeting siRNAs (NT) or siRNA targeting PQBP1, were challenged with HIV-1 for 2.5 hours and stained for cGAMP (green), cGAS (red), and dapi (blue). Mean cGAMP signal per cell were graphed (bottom). Median and Error bar (-/+ 1.5*IQR) are shown. Box indicates the interquartile range (IQR). Mann-Whitney **** p<0.0001. Data are representative of three independent infections. All the data presented, unless otherwise stated, are representatives of at least two independent experiments.

The temporal disparity between PQBP1 and cGAS recruitment to the capsid suggests a mechanism wherein PQBP1 and capsid association is a prerequisite for cGAS recruitment. These results also suggest that binding of PQBP1 to capsid is not sufficient to recruit cGAS, but that loss of capsid integrity associated with reverse transcription progression licenses PQBP1 recruitment of cGAS to the HIV complexes. Since genetic ablation of PQBP1 is incompatible with cell viability (Iwasaki and Thomsen, 2014; Tamura et al., 2013; Yoh et al., 2015), we transiently depleted PQBP1 from THP-1 cells, and evaluated the frequency of cGAS association with the virus particles (Figure 3C). Reduction of PQBP1 protein in cells was confirmed by a decrease in both PQBP1 mRNA and its protein level (Figure S3C, top). Importantly, down-regulation of PQBP1 resulted in a marked decrease in cGAS association per virus particle (Figure 3C), as well as a marked drop in the fraction of virus particles that are targeted by cGAS (Figure 3D) without a significant overall reduction of cGAS protein levels (Figure S3C bottom). Consistent with a model wherein cGAS is specifically concentrated to PQBP1-decorated virions, we observed that a significantly higher density of cGAS foci is present in the vicinity of virions when compared to the randomly distributed ones in the cytoplasm (data not shown). Lastly, we addressed the contribution of the PQBP1/cGAS/capsid complex formation for the innate immune response to HIV-1 infection by assessing cGAS activation through visualization of cGAMP levels (green) in cells (Hall et al., 2017). THP-1 cells, either infected with HIV-1 or transfected with HT-DNA for 3 hours followed by immunostaining, revealed cGAMP staining in the challenged cells that are analogous to the ones where cGAMP molecules were transfected (Figure S3D). Consistent with a robust induction of cGAMP, we also observed a nuclear localization of IRF3, a hallmark of the cGAS/IRF3 pathway activation, in the challenged cells (IRF3 panel, Figure S3D). Accordingly, a selective depletion of PQBP1 resulted in a marked decrease in the cGAMP level in response to HIV-1 challenge, reconfirming the PQBP1 dependency for the cGAS activation (Figures 3D and S3E).

Previously, Lahaye et. al. have shown that NONO associates with nuclear HIV capsids and is needed for innate immune responses upon HIV infection in MDDCs (Lahaye et al., 2018). Consistent with results reporterd by Lahaye et. al., we observe HIV-1 infection induces cGAS signaling in a NONO-dependent manner in MDDCs (Figures 4A and S4A). However, we find that NONO depletion in THP-1 cells, which maintain an intact innate response to HIV (Collins et al., 2015; Gao et al., 2013; Sumner et al., 2020; Sun et al., 2013; Wiser et al., 2020; Yoh et al., 2015), has no significant effect on innate immune response to HIV-1(with Vpx), as well as HIV-2, infection (Figures 4B). In attempt to determine whether NONO participates in the initial recognition of incoming virus particles in MDDCs, we evaluated the frequency of NONO association with incoming HIV-1 virions. The distributions of nearest neighbor distance analysis revealed that IN and NONO association displayed a single broad peak at ∼0.9 µm (blue lines, Top) that is similar to the distribution of the random curve generated upon the *in silico* shuffling of the foci coordinates (magenta, Top, Figure 4C). In contrast, IN to PQBP1 distance distribution showed a distinctive peak at < ∼0.5 µm (blue lines, Bottom) which represents INs that are colocalizing with PQBP1, separating away from the randomized shuffling of the coordinates (magenta, Bottom, Figure 4C; see Figure 1 also). At a threshold of 0.5 µm, we find that ∼ 8 % of INs associate with NONO (similar to the level of randomized control) while ∼ 23% of INs do with PQBP1 (Bottom left, Figure 4C). These data indicate that NONO does not significantly colocalize with incoming virus particles (INs) as PQBP1 does in MDDCs.

**Figure 4.**
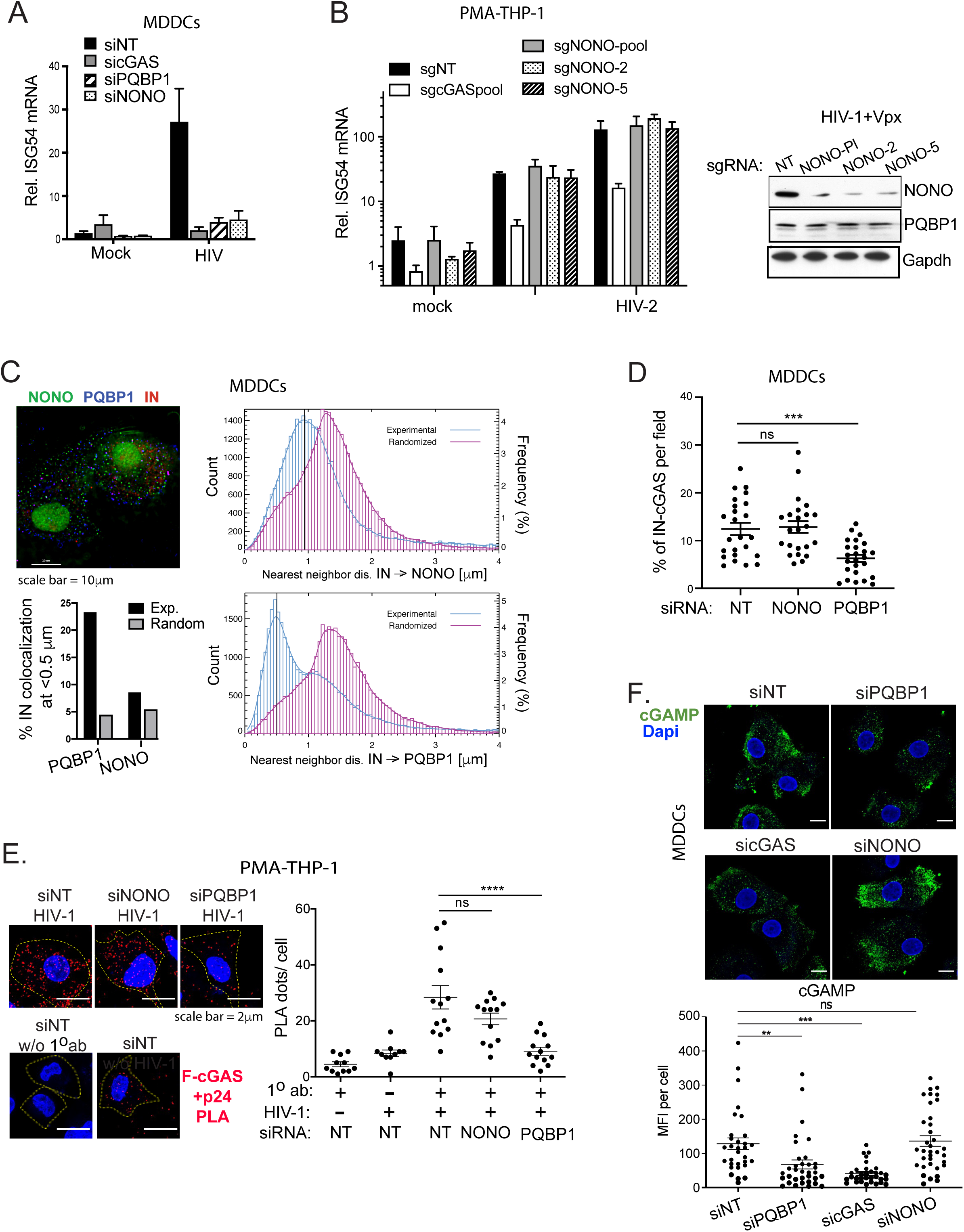
NONO is not required for the PQBP1-dependent cGAS sensing occurring during the early step of the infection. (A) NONO is required for the HIV-1 infection induced ISG54 induction in MDDCs. Cells targeted by the indicated siRNAs were either mock-treated or infected with HIV-1 luciferase virus, followed by ISG54 mRNA measurement 16 hrs post infection. (B) NONO is not required for ISG54 induction against HIV-1 infection in PMA differentiated THP-1 cells. Left, Cells were subjected to CRISPR gene editing with indicated sgRNAs and assessed for ISG54 mRNA inductions at 16 hours post infection with either mock, HIV-1 in the presence of VLP-Vpx or HIV-2. Right, the levels of NONO and PQBP1 proteins of the cells targeted with indicated sgRNAs were shown accordingly. (C) Left, representative image of MDDCs infected with HIV-1 virus (IN-mRuby3) and stained for PQBP1 (blue) and NONO (green). Right, distribution of IN-to-NONO (top) or IN-to-PQBP1 (bottom) nearest neighbor distances (*d*). The distribution of experimental data (blue histograms) is compared with one generated from *in silico* randomized dots (magenta histograms). A kernel density estimate of each distribution is overplotted as a solid curve, represented as Frequency, N=27,932 INs. The percentage of INs that have either PQBP1 or NONO at *d* < 0.5 µm as well as the values for the randomized controls are graphed bottom left. (D) NONO depletion does not impair cGAS recruitment to incoming capsids. MDDCs treated with indicated siRNAs were infected with HIV-1 viruses (Gag-IN-mRuby3/GFP) for 2.5 hours, followed by immunostaining for cGAS and imaging. Fractions of total IN foci overlap with cGAS signal are graphed. Mean and SEM are shown. One-way ANOVA, *** p<0.001. The results are based on and four independent experiments. (E) NONO is not required for cGAS recruitment to capsid in PMA-THP-1 cells. Efficiency of Flag-cGAS and p24 viral capsid interaction in PMA-THP-1 cells targeted by indicated siRNAs was determined by a proximal ligation assay (PLA). The cells were infected with HIV-1 virus in the presence of VLP-Vpx for 2 hrs followed by paraformaldehyde fixation and PLA. Representative images (left) and quantification (right) of PLA dots (red) per cells are shown. Dapi (blue) and cell boundary (dotted line) are shown. Mean and SEM are shown. One-way ANOVA, **** p<0.0001. ns denotes no significance. (F) cGAMP production upon HIV-1 infection is not impaired by depletion of NONO. MDDCs, transfected with indicated siRNAs, were challenged with HIV-1 for 2.5 hours and were stained for cGAMP (green) and dapi (blue). Mean cGAMP signal per cell were graphed (bottom). Mean and SEM are shown. One-way ANOVA, ** p<0.01, *** p<0.001, ns denotes no significance. All the data, unless noted otherwise, are representative of at least two independent experiments.

Next, we examined whether NONO is required for cGAS recruitment to incoming virus particles. Analogous to Figure 3A, the puncta overlap between IN-mRuby3 (virus) and cGAS signals were assessed in the infected MDDCs and determined % of total INs positive of cGAS signal. We observed that depletion of NONO had no noticeable impact on cGAS co-association with INs (Figure 4D; Figure S4B). Similarly, we find that NONO depletion did not impact cGAS association with the viral capsid in THP-1 cells as well. Specifically, a proximal ligation assay between Flag-cGAS, stably expressing, and p24 CA in the infected THP-1 cells confirms that NONO is dispensable for the cGAS recruitment to incoming capsids (Figure 4E; Figure S4C). Lastly, we confirmed that loss of NONO did not impact cGAMP production in MDDCs upon HIV- 1 (Figure 4F; Figure S5C). Taken together, these data suggest that, in certain cellular contexts, NONO is dispensable for the innate response to HIV. Moreover, these results also indicate that NONO does not participate in the recruitment of cGAS to incoming HIV-1 capsid and its subsequent enzymatic activation.

### PQBP1 interaction with capsid licenses cGAS sensing of HIV-1 infection

Upon observation of PQBP1/cGAS complex assembly on HIV-1 capsid, we asked whether the capsid could serve as a platform where PQBP1 and cGAS interaction can be facilitated. Immunofluorescence (IF) based colocalization analysis revealed that HIV-1 infection selectively enhanced PQBP1 and cGAS co-association whereas a direct activation of cGAS by herring-testis (HT) DNA did not (Figure 5A). Briefly, the colocalization analysis was determined with three-dimensional detection of spots, performing image segmentation. We utilized a threshold distance of 0.4 μm to define the interaction between both molecules. We observed a seven-fold increase in the frequency of PQBP1-cGAS colocalization for HIV-1 infected cells, while no enhancement in association was observed in cells transfected with HT-DNA, an established cGAS ligand (right, Figure 5A). Complementing the IF result, we also demonstrated that PQBP1 co-precipitated with cGAS more efficiently in the presence of HIV-1 infection but not in the HT-DNA treatment compared to the no treatment control of both MDDCs and PMA-THP-1 cells (Figure 5B). A systematic truncation analysis on PQBP1 protein confirmed that cGAS interaction surface is distinctive from the capsid interaction domain (Figure S5A). Briefly, either full-length or truncated PQBP1-YFP proteins was over-expressed with MBP-tagged cGAS proteins in 293T cells and subjected to co-immunoprecipitation assays (coIPs). A mutant PQBP1 that lacked the capsid binding domain of first 46 amino acids (aa 47-265) co-precipitated with MBP-cGAS as efficiently as the full-length (aa 1-265), yet the capsid-associating N-terminal domains of PQBP1 (aa 1-46, as well as aa 1-104) failed to do so (Figure S5A). Previously we have shown that a point mutation with in the WW domain (aa 47-86) of PQBP1 impair cGAS interaction; however, we observed that the N-terminal fragment containing WW domain (aa 1-104) is not sufficient but need additional c-terminal regions to bind to cGAS [compare aa 1-104 with aa 1-211 in Figure S5; (Yoh et al., 2015)]. Collectively, the reciprocal result of the coIP suggests that two distinctive interaction surfaces exist within PQBP1, one for capsid and the other for cGAS interaction.

**Figure 5.**
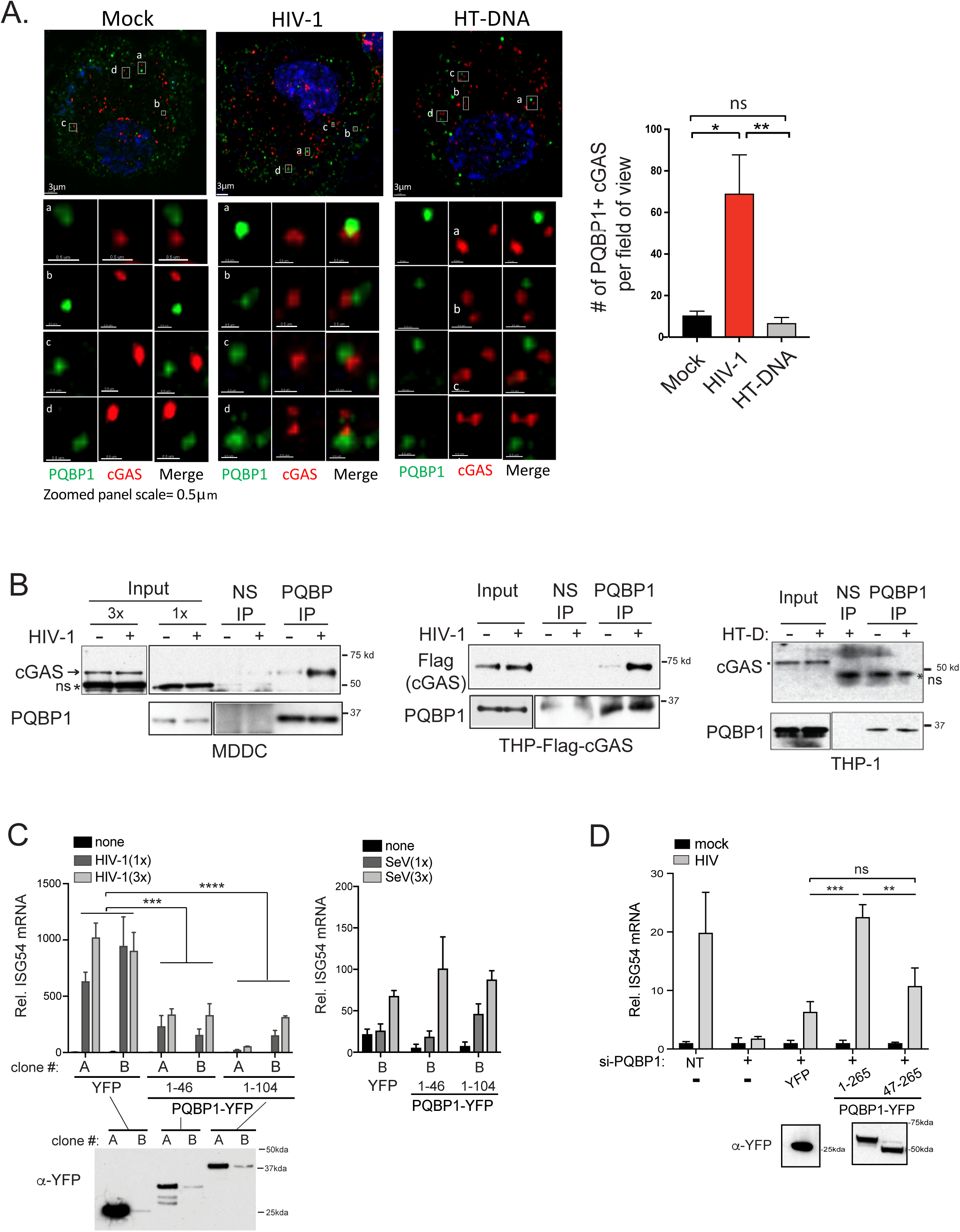
HIV-1 infection promotes the formation of PQBP1/cGAS innate sensing complex. HIV-1 infection enhances PQBP1 and cGAS interaction. (A) PMA-THP-1 cells were fixed 2.5 hours post HIV-1 infection (Gag-IN-mRuby) or HT DNA transfection, followed by immunostaining against PQBP1 and cGAS and for confocal microscopy. Z-section images (left) and the quantification of the co-association foci (right) are shown. A threshold for colocalization is set at a distance of 0.4 µm. Averages and SEM are shown. One-way ANOVA, ** p<0.01, * p<0.05, ns denotes no significance. (B) HIV-1 infection (+) enhances co-immunoprecipitation of cGAS or Flag-cGAS with PQBP-IP in MDDCs (left) or PMA-THP-1 (middle and right), respectively. The cells were infected with HIV-1 luciferase virus in the presence of VLP-Vpx and subjected to co- IPs at 3 hrs post infection. HT-DNA was delivered to PMA-THP-1 cells instead of the viral infection and endogenous cGAS co-precipitating with PQBP1 was assayed. 1x and 3 x denote the relative amounts of inputs loaded on a gel. Normal IgG (NS) or antibody against PQBP1 were used as indicated. ns denotes non-specific bands. (C) The capsid interaction domain of PQBP1 is a dominant inhibitor of cGAS-mediated innate sensing of HIV-1 infection. Two independent PMA- THP-1 cell clones (A and B clones) stably expressing either YFP, PQBP1__1-46-_YFP or PQBP1__1-104-_YFP were infected with HIV-1 luciferase virus in the presence of VLP-Vpx or Sendai virus as indicated. ISG54 mRNAs were measured 16 hours post infection. (D) The capsid interaction domain of PQBP1 is needed for innate response against HIV-1 infection. PMA-THP-1 cells either un-transduced or stably expressing the indicated PQBP1-YFP proteins were treated with siRNA against endogenous PQBP1 (+), followed by HIV-1 infection and ISG54 mRNA detection as in (C). NT denotes non-targeting siRNA. ISG54 mRNA levels were expressed as a fold induction over their uninfected counterparts. Expression level of either YFP or PQBP-YFPs in the cells used in (C) and (D) were determined by anti-GFP/YFP western blots. Equal numbers of cells were analyzed. T-test (unpaired; two-tailed) *p<0.05, **p<0.01, ***p<0.001. All the data, unless noted otherwise, are representative of at least two independent experiments.

We found that the first 46 a.a. of PQBP1, which excludes cGAS binding surface, is sufficient to bind to capsid in cells (Figure 2D-E). We next asked whether this domain possesses a dominant negative impact on cGAS sensing of HIV-1 infection by potentially competing off endogenous PQBP1 bound to capsid, and in turn reducing cGAS recruitment to the virus particles. Ectopic expressions of 1-46 and 1-104 aa of PQBP1 inhibited ISG54 mRNA induction to HIV-1 infection compared to the responses from the cells expressing YFP alone (left, Figure 5C). This observed effect was specific to HIV-1 infection, since the inhibition was not apparent with Sendai virus infection (right, Figure 5C). Reciprocally, THP-1 cells expressing a mutant PQBP1 that lacked the capsid binding surface (residues 47-265) but retained cGAS interaction (Figure S6A) were impaired in ISG54 mRNA induction in response to HIV-1 challenge (Figure 5D), underscoring the importance of PQBP1 and capsid interaction for cGAS recruitment and innate sensing of HIV-1 infection.

## DISCUSSION

In this study, we find that the assembled capsid lattice acts as a preliminary PAMP that serves to authenticate incoming viral nucleic acid cargo. Specifically, we find that PQBP1 recognition of the multimerized capsid lattice, likely at the positively charged arginine ring in the CA, acts as an initial PAMP engagement step that initiates an innate immune program. Consistent with this finding, we do not observe significant interaction between PQBP1 and monomeric CA proteins (data not shown). Importantly, the charged pore structure is highly conserved among the capsids of the lentivirus family (Mallery et al., 2019), which is consistent with our previous observation where PQBP1 is a lentivirus specific co-factor for cGAS sensing of viral DNA. The intact capsid cone has estimated ∼250 CA protein hexamers plus twelve pentamers that would provide numerous binding sites for the accumulation of PQBP1 molecules (Perilla and Gronenborn, 2016; Pornillos et al., 2011). Indeed, *in vitro* binding assays indicated that multiple PQBP1 proteins are bound to each assembled CA lattice (compare AF488-PQBP1 signal peaks in the absence and presence of CA-A204C, Figure 2B). Through its heterodimerization with PQBP1-bound capsid, cGAS becomes co-positioned at the site of PAMP generation. This recruitment functions to license the second step of the innate immune recognition of lentiviral infection: induction of a vigorous cGAS-dependent response. This dual-step recognition of both pathogen protein and DNA likely allows for pathogen sensitivity and specificity to ensure that the activation of IFN pathways is not spontaneously activated from recognition of each individual PAMP separately, assuring a powerful yet specific inflammatory response.

The prerequisite of PQBP1 binding to capsid suggests that additional events are required for cGAS recruitment to incoming viral particles. The finding that cGAS is recruited to the PQBP1- capsid platform only upon the initiation of capsid disassembly (Figure 3B) suggests that molecular events that may be coupled to disassembly are required for the formation of a competent PRR complex. These may include conformational changes in PQBP1, oligomerization/activation of cGAS, progression of reverse-transcription and/or the cytosolic exposure of reverse transcribed DNA upon the loss of the core integrity (Christensen et al., 2020; Mamede et al., 2017; Manel et al., 2010; Sumner et al., 2020). Insights into the potential nature of the RTC, PQBP1, and cGAS complex come from various studies. The Inhibition of reverse transcription by nevirapine treatment blocked both capsid structural integrity loss and innate response (Felts et al., 2011; Ilina et al., 2012; Mamede et al., 2017; Wang et al., 2021; Yoh et al., 2015). Blocking of the first strand transfer of the reverse transcription with a RNase H inhibitor (Ilina et al., 2012; Julias et al., 2002) did not compromise the innate response (data not shown), which suggests that the initiation of reverse transcription, i.e. production of the strong stop HIV-1 DNA is sufficient to induce cGAS activation. The coupling between the reverse transcription and capsid disassembly is solidified by an identification of capsid mutant virus with an accelerated reverse transcription kinetic displaying faster initiation of capsid disassembly (Sultana et al., 2019). Lastly, cryo-electron tomograph analysis of the *in vitro* assembled reverse transcription complexes reveals strand-like loops, believed to be the reverse transcribing DNA extruding out from a disassembling capsid (Christensen et al., 2020). Collectively, these data underscore the crosstalk among reverse transcription, capsid disassembly and cGAS mediated innate sensing. Whether the initial structural changes in capsid triggered by the progression of reverse transcription permits the recruitment and activation of cGAS, or the availability of the reverse transcribed DNA providing an additional tether for the process, will need to be investigated.

Lahaye et. al. have reported that cGAS sensing of HIV can occur nucleus through binding of disassembled capsid (Lahaye et al., 2018). In contrast, we have shown that the recognition of intact capsid of incoming virions in the cytoplasm by the PQBP1/cGAS complex is required for establishing the innate response to HIV-1, and occurs independently of NONO. These observations raise the possibility that cytoplasmic and nuclear sensing of HIV may be functionally coupled through capsid, but PQBP1 detection of incoming HIV-1 is a prerequisite for PAMP authentication and initiation of innate immune response to viral challenge.

Collectively, these data reveal a unique two-factor authentication strategy that enables cellular immune response to transient and low-abundance retroviral DNA species. PQBP1 binding to HIV-1 capsid is a component of a multi-step sensing process that enables cGAS recruitment and DNA PAMP recognition, culminating in the initiation of an innate immune response against HIV-1 infection. This modular association of PQBP1 with the cGAS pattern recognition receptor triggers a robust response to veritable foreign DNA species, while circumventing self-activation from extranuclear host-derived DNA. This molecular strategy represents an evolutionary expedient mechanism to expand the versatility of germline-encoded sensors to mount effective immune responses across a range of invading pathogens with unique features that associate them as PAMPs.

## ACKNOWLEDGEMENTS

The authors would like to thank Zeli Zhang for reagent preparations; SBP Flow Cytometry core for technical assistance.

## AUTHOR CONTRIBUTIONS

Conceptualization, S.M.Y., J.I.M. and S.K.C; Methodology, S.M.Y., J.I.M. and D.L.; Validation, A.T.; Formal Analysis, G.C.C., L.R., J.I.M.; Investigation, S.M.Y., J.I.M., D.L., N.A., M.T. S-A., J.T., N.H-F.; Resources, H.C., L.S., S.G, J.H.; Writing-Original Draft, S.M.Y, J.I.M., D.L., T.B.; Writing-Review & Editing, S.K.C., A.G-S., T.H., Y.J., R.K.; Visualization, S.M.Y., J.I.M., T.B.; Supervision, S.K.C., T.H., A.G-S.; Project Administration, S.M.Y.; Funding Acquisition, S.K.C., A.G-S., S.M.Y.

## DATA AND MATERIAL AVAILABILITY

All data, code, and materials used in this work are available in the main text or the supplementary materials and upon request.

### STAR METHODS

#### Reagents

50 – 100 ng of cGAMP (Invivogen) and 2 ng to 10 ng of Herring testis (HT) DNA (Sigma) were transfected into 2.5 × 10^4^ PMA-differentiated THP-1 using Lipofectamine 2000 (Life Technologies). The following antibodies were used: IRF3 (Cell Signaling D9J5Q), PQBP1 [Bethyl Laboratory A302-802A; Santa Cruz Biotechnology sc-376039; Sigma Aldrich (1A11)], cGAS [Novus NBP1- 8676; Santa Cruz Biotechnology (D-9); Cayman (5G10); Cell signaling (D1D3G)], normal rabbit/mouse IgG (Santa Cruz Biotechnology SC-2027), β-actin (Cell Signaling Technology 49705), GFP (Thermo Scientific MA5-15256: Clontech 632592), FLAG (Sigma Aldrich F1804), p24 (AIDS Reagent 71-31). Anti-cGAMP [PF-07043030 Pfizer (Hall et al., 2017)], Anti-MBP magnetic beads (NEB).

#### Plasmids, siRNAs and qRT-PCR

Human LentiORF cDNA clones of PQBP1 (Open Biosystems), both wild-type and mutant constructs, in-frame fused with eYFP proteins, were generated by a standard Gibson cloning approach, and packaged into lentiviruses according to the manufacturer’s protocol. The silent mutations were introduced to PQBP1-eYFP constructs to render the proteins resistant to siPQBP1 RNAs as described previously (Yoh et al., 2015). Flag-cGAS is cloned into pEASIL doxycline inducible lentivirus vector (a generous gift from M. Malim). MBP-cGAS and 6xHIS- PQBP1 constructs were cloned into pCDNA 3.1 and pCDFDuet vectors respectively. THP-1 cells, either wildtype or Dual-KO-cGAS (Invivogen) were infected with each lentivirus to generate the cells with a stable expression of the protein of interest.

siRNAs were introduced into PMA-THP-1 cells using Stemfect RNA transfection kits. Typically, 5 pmol of siRNAs and 0.17 µl of Stemfect were used for 2.5 × 10^4^ cells. Forty-eight hour after siRNA transfection, the cells were infected with either HIV-1 virus in the presence of VLP- VPX (Manel et al., 2010; Yoh et al., 2015) or 2.5 HAU/ml of Sendai virus for 16 hours, followed by RNA isolation and qRT-PCR analysis. Alternatively, in the case of IF analysis, cells were fixed 2-3 hours post infection. Following siRNAs were used: siNT (5’-AATCGATCATAGGACGAACGC- 3’); siPQBP1-1 (5’- AAGCTCAGAAGCAGTAATGCA-3’); siPQBP1-2 (5’-AAAGCCATGACAAGTCGGACA-3’). qPCR primers that were used were previously described (Yoh et al., 2015).

#### Cell culture and viral infection

This study was approved by the National Institutes of Health (NIH) through our Institute Biosafety Committee (IBC). Primary monocyte-derived dendritic cells (MDDCs) were prepared from fresh, healthy donor blood from the San Diego Blood Bank as described previously (Yoh et al., 2015). The human monocyte-like THP-1 cell line, grown in RPMI 1640 supplemented with 10% FBS, was differentiated by treating with 20-40 ng/ml PMA (Phorbol myristate acetate) for 2 days. Both THP-1 and HEK293T cells were purchased from ATCC. THP-1 Dual and THP-1-Dual KO-cGAS cells were purchased from Invivogen and maintained in RPMI 1640+10% FBS.

MDDCs or PMA-differentiated THP-1 cells were infected with VSV-G pseudo-typed HIV- 1 virus and VLP-Vpx, as described previously (Manel et al., 2010; Yoh et al., 2015) unless otherwise specified. In general, 2-10 ng of p24 or 0.1 to 0.5 RT unit of HIV-1 viruses were used per 25K cells in 100 μl of media and harvested between 1.5 to 16 hours post infection as specified. All HIV-1 viruses are generated by transient transfection of provirus plasmids in 293T in 10cm dish at 50% confluency and harvesting 48 hours post transfection, followed by DNase-treatment and concentrated through a 25% sucrose TNE by centrifugation for 16 hours at 3000 × g at 4°C. VSV-G-pseudotyped HIV-1 firefly luciferase (luc) was prepared by transfecting 10 µg of NL4-3 R+ E- firefly-luc plasmid, provided by Dr. Nathaniel Landau and 1.5 μg of pCMV-VSV-G with 40 µl of PEI (pH 4.5, 1 µg/ml) or 30 µl of lipofectamine 2000. HIV-Gag-IN-mRuby3 virus were produced by transfecting 5 μg of pNL43 env-, 2 μg of Gag-IN-mRuby3 and 4 μg of pCMV-VSV-G. HIV- iGFP and Gag-IN-mRuby3 viruses were produced by transfection of 5 μg of HIV-Gag-iGFP, 3 μg of pGag-IN-mRuby3, and 4 μg of pCMV–VSV-G (Mamede et al., 2017; Sultana et al., 2019). CypA-dsRed packaged HIV-1 virus were produced by transfection of 5 μg of NL43dEnv-, 3 μg of pCMV-VSV-G and 4 μg of CypA-dsRed plasmids (Francis et al., 2016; Francis and Melikyan, 2018).

The Mount Sinai Department of Microbiology Virus Collection provided the Cantell strain of Sendai virus, and it was grown for 2 days in 10-day old embryonated chicken eggs, and it was tittered using turkey RBC HA assays (Lampire Biological Laboratories).

#### CRISPR-Cas9 RNP Production and THP-1 Electroporation

Detailed protocols for RNP production have been previously published (Hultquist et al., 2019). All crRNA guide sequences used in this study were derived from the Dharmacon pre-designed Edit- R library for gene knock-out, including the non-targeting guide (U-007502), cGAS-targeting guides (equimolar pool of CM-015607-01 through CM-015607-05), and NONO-targeting guides (CM- 007756-02 and CM-007756-05).

#### Guide RNA Sequences

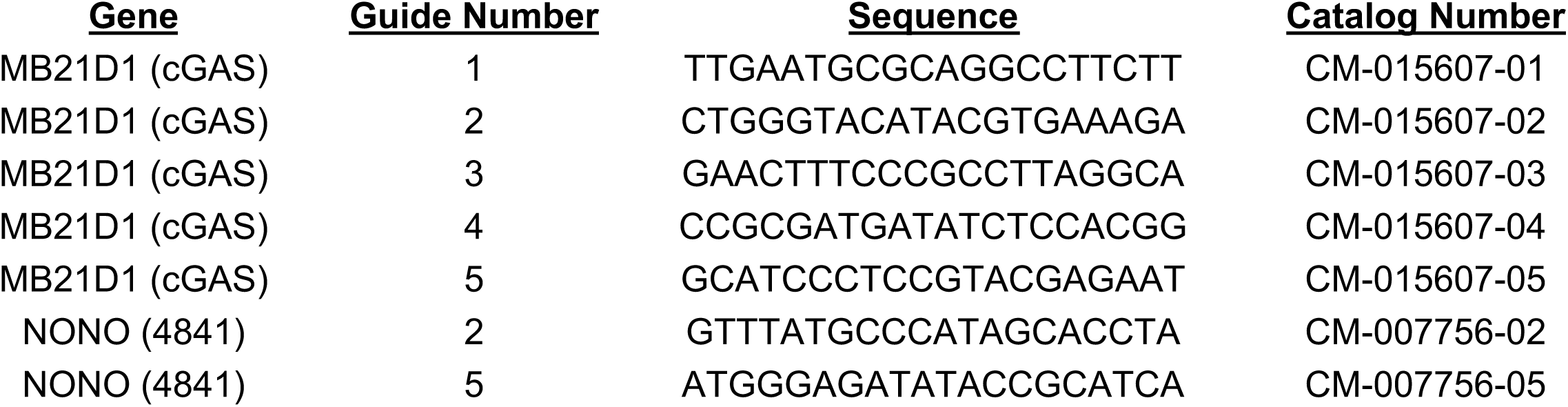

#### Immunofluorescence, image acquisition and analysis

PMA-THP-1 cells were seeded at a density of 2.5 × 10^4^ cells/ well in a 96 well glass plate for 48 hours followed by infection with HIV-1 virus with or without VLP-Vpx as specified or transfection of either HT DNA or cGAMP for 1.5 hrs to 3 hrs. The cells were then fixed with 4% w/v paraformaldehyde in PBS and permeabilized using a standard protocol and subjected to immunostaining. Cells were imaged via fluorescence wide-field deconvolution or confocal microscopy. DV-Elite, GE-Ultra, OMX-SR, Nikon TIe-2, Zeiss LSMv880 were utilized. Images obtained in GE microscopes were deconvolved in SoftWorx package with the standard vendor software definitions and camera biases. Images obtained in Nikon TIe-2 microscope were deconvolved using FlowDec using Lucy-Richardson algorithm with the support of pims using python 3.x with PSFs generated in FIJI/ImageJ. Super resolution Images were obtained in 3D- SIM mode in a GE OMX-SR microscope equipped with 4 laser lines, 60x/1.42 NA oil immersion lens (Olympus), and 3 independent cameras. Images were calculated/reconstructed and channel registered via the vendor’s software (SoftWorx) with 0.001 Wiener constant values.

#### Fluorescence resonance energy transfer assay

Viral infections with dsRed-CypA labeled particles were imaged in cells expressing eYFP or PQBP1-eYFP truncations. Images were acquired as a Z-stack with a GE-Deltavision Ultra microscope equipped with a PCO edge CMOS camera and an oil immersion 60x/1.42 NA Olympus lens with 1.42 NA and deconvolved using the vendors software (SoftWorkx). Excitation was done with an SSI-LED light source, and the emitted wavelength collection was cleaned up with the different excitation/emission filters; for FRET (YFP/mCherry); YFP (YFP/YFP); dsRed (mCherry/mCherry). Analysis was performed by identifying the coordinates of viral particles in the deconvolved images and measuring the intensities from the raw images on each corresponding channels in the regions containing viral particles. Cross-talk was controlled by measuring intensities of dsRed-CypA labeled particles in cells in the absence and the presence of eYFP or PQBP1-eYFPs constructs. The cross-talk control also included the measurement of signals form the cells without viral particles. The values for the normalized FRET of viral particle Intensities were calculated using following formula and methods as in (Jiang and Sorkin, 2002; Sood et al., 2017; Zal and Gascoigne, 2004):

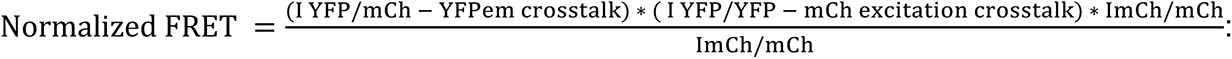

#### Viral capsid integrity assay - Correlative Live and fixed immunofluorescent Imaging Analysis

Delta Vision wide-field microscopes (DV-ELITE or OMX-SR, GE Life Sciences) equipped with an electron multiplying charge-coupled device (EMCCD) camera and CMOS cameras (PCO edge), solid state illumination (SSI-LED) light path was used to acquire time-lapsed fluorescent snapshots HIV-iGFP/IN-mRuby3 viruses infecting THP-1 or MDDCs plated in Delta T culture dishes (Bioptechs) or μ-Slide I Luer (Ibidi) that were coated with fibronectin (SIGMA) for overnight adhesion of MDDCs (but not THP-1) according to the supplier’s instructions. Cells were kept in a 37°C heated chamber, together with a blood gas mixture (5% CO_2_, 20% oxygen), throughout the imaging process. Cells were incubated with RPMI without phenol red with 10% FBS, L-glutamine, and MEM-NEAA. All infections were done with Polybrene at a concentration of 5 μg/ml. Z-stacking spacing was set to 0.5 µm, with a total of 12-μm z-axis imaging for the fluorescence snapshots, and a single Z reference image was taken in bright field for cell edge identification. Nominal magnification was 60x/1.42NA lens for all experiments. Cells were washed once with PBS, promptly after the last time point of imaging, and fixed with final 3.7% formaldehyde in piperazine-N,N′-bis (PIPES) buffer for 5 min, followed by three PBS washes, as in (Mamede et al., 2017). Culture dishes were then permeabilized with blocking media made of TX-10 donkey serum and stained with antibodies probing for PQBP1 (Sigma) and cGAS (Novus) followed by anti-mouse- AF647 and anti-Rb-dylight405 (Jackson Immunoresearch) secondary antibodies, respectively. Using the same microscope, or synchronized trays between DV-ELITE and OMX-SR, the previously time-lapse imaged cells and fields of view were found and imaged after staining. The viral particles were identified by their IN-mRuby3 signal present in the same area as the last time- lapse time-frame, when still present in this second phase. ROIs were set in ImageJ/Fiji based on the IN-mRuby3 puncta and the mean intensities for all channels in such ROI were quantified and plotted. The second verification of capsid integrity loss was performed in the fixed imaging stage by the disappearance of iGFP signal.

Z-stacks were deconvolved and z-projected using SoftWorx (GE Life Sciences) before each individual IN-mRuby3 particle were tracked over time was performed using FIJI/ImageJ (NIH). Mean intensities of HIV-iGFP were automatically measured in the same x-y coordinates where the IN particle was identified by the tracking algorithm. Centered particle video recordings were automatically generated by an in-house–made Python scripts using pims (http://soft-matter.github.io/pims/) and Matplotlib libraries, with the data analyzed and exported from Fiji/ImageJ as in (Mamede et al., 2017; Sultana et al., 2019).

##### Proximal Ligation Assay (PLA)

PMA-THP-1 cells, 3 hours post-infection, were fixed in 4% PFA in PBS for 20 min, permeabilized with 0.2% Triton X-100 in PBS, and subjected to PLA assay according to manufactural protocol of Duolink in situ detection reagents red kit (DUO92008, Sigma-Aldrich). Antibody imcubations were used at 1;400 dilution: anti-rabbit HIV1 p24 (Abcam- ab32352) and mouse anti-FLAG® M2 antibody (Sigma). PLA dots were detected using Nikon A1R HD confocal with 60X objective. Analysis was done using ImageJ.

##### Image Analysis

Identification of viral particles from infected THP-1 cells was done with python with skimage, scipy, pims, and trackpy tools (v0.4.2 - http://soft-matter.github.io/trackpy/). In short, cells were subjected to a binary threshold so that no extracellular signals (viruses or cell debris) would confound the analysis. Viral particles were detected using thresholding and watershed segmentation methods or by using trackpy. For the virus analysis, the masks of the pixels that are positive for viral particles were then quantified for each independent channel (PQBP1, cGAS, iGFP, IN label, etc)

#### Co-immunoprecipitation

MDDCs or PMA-THP-1 cells expressing Flag-cGAS proteins were infected with VSV-G pseudo- typed HIV-1 luciferase virus at MOI of 1 in the absence and presence of VLP-Vpx co-infection respectively. Three hours post infection, cells were lysed and subjected to immunoprecipitation against endogenous PQBP1 proteins, followed probing the IPs for endogenous cGAS or Flag- cGAS presence. HEK293T lysates expressing eYFP or eYFP fused either to wild type or mutant PQBP1 and MBP-cGAS were subjected to immunoprecipitation against MBP tags, followed by probing for the presence of PQBP1-YFP proteins. Typically, 300 μg of protein lysates were incubated with anti-MBP-magnetic beads (NEB) in 100 mM KCl, 12.5mM MgCl2, 0.5% Triton-X, 20 mM HEPES, 0.2 mM EDTA, 10% glycerol, 0.2 μM PMSF, and protease inhibitors. The beads were washed three times with a washing buffer (20 mM HEPES, 0.2 mM EDTA, 300 mM KCl, 0.5% Triton-X, 10% glycerol and 0.2 μM PMSF) followed by a conventional western blot analysis.

#### Recombinant protein production

C-terminal MBP-6xHis tag is fused to CA (A14C/E45C) using a SARS main protease cleavage site linker in pET11a as previous described (Summers et al., 2019). 6xHis-PQBP1 was cloned into pCDFDuet using BamHI/HindIII restriction sites. MBP, used as a negative control in co- pelleting, was obtained from cleaved CA(A14C/E45C) during purification before assembling into tubes. Protein was over expressed in BL21(DE3) cells grown to OD of 0.6-0.8 and induced with 0.5 mM IPTG at 25 °C for CA(A14C/E45C)-MBP-6xHis or 18°C for 6xHis-PQBP1 for 18 hours. Cells were lysed via microfluidization in 50 mM Tris, pH 8.0, 500 mM NaCl, 5% v/v glycerol, 0.1 mM TCEP with a protease inhibitor tablet (Roche). CA(A14C/E45C)-MBP-6xHis was purified as described previously (Summers et al., 2019). 6xHis-PQBP1 clarified lysate was purified via Ni- NTA (Qiagen) affinity and size exclusion chromatography (GE). 6xHis-PQBP1 was concentrated to 5-10 mg/mL in 50 mM Tris, pH 8.0, 50 mM NaCl, 0.1 mM TCEP and flash frozen in liquid nitrogen until use. CA tubes were assembled as described previously (Pornillos et al., 2009) from the purified, MBP-6xHis cleaved CA(A14C/E45C) and stored at 4°C in 50 mM Tris, pH 8.0 until use.

Co-pelleting assays were performed by incubating a final concentration of 10 µM PQBP1 or MBP with 100 µM assembled CA tubes in the presence of 25 or 500 mM NaCl on ice. Reaction buffer contained the noted amount of NaCl with 50 mM Tris, pH 8.0, 25 mM NaCl, 0.1% v/v NP-40. Samples were centrifuged 20 min at 4°C and 14,500 g and supernatant was separated from the pellet. The pellet was dissolved in an equal volume of buffer and samples were analyzed via SDS-PAGE on NuPage™ gel (Invitrogen) and developed with SimplyBlue™ stain (Invitrogen). Gel was imaged directly with a CCD camera.

#### Covalent labelling of Recombinant PQBP1, CypA, and CPSF6_313-327_

Recombinant PQBP1 proteins in PBS (1X) and 0.1 mM TCEP was incubated with AlexaFluor488- C5-malamide dye (Thermo Fischer Scientific) in a 1:1.2 molar ratio for 15 min at room temperature. Unreacted dye was removed by size exclusion chromatography using a Superdex 200 Increase 5/150 GL column (Cytiva) equilibrated in 20 mM Tris-HCl (pH 8.0) and 100 mM NaCl flowing at 0.1 mL/min on a HPLC (Shimadzu). Fractions were collected manually and fractions containing labelled AF488-PQBP1 conjugate were identified by SDS PAGE and imaging with the Alexa Fluor 488 filter on the ChemiDoc MP imaging system (BIORAD). Proteins were stored at -80°C in 10% v/v glycerol and dialyzed into 20 mM Tris-HCl (pH 8.0) and 75 mM NaCl prior to TCCD measurements. Purification and labelling of CypA and CPSF6_313-327_ peptide with Alexa Fluor dyes were previously described (Lau et al., 2019; Peng et al., 2019).

#### Cell-Free Expression and Purification of GFP-tagged PQBP1 fragments

The coding sequences for PQBP1_1-46_ and PQBP_147-265_ were amplified by PCR and cloned into Gateway™ vectors and contain either an N-terminal 8xHis-eGFP or C-terminal sfGFP-8xHis tags using Gibson assembly (NEB). For each construct, purified plasmid DNA was added to 100 µL *Leishmania tarentolae* cell-free expression mix to 60 nM DNA and expressed for 2.5 hrs at 28 °C.(Lau et al., 2019) The expressed PQBP1 fragments were bound to Ni-NTA beads (BIORAD) for 30 min on ice and washed with 300 µL of 20 mM Tris-HCl (pH 8.0) and 100 mM NaCl before eluting bound protein with 40 µL of the same buffer containing 0.5 M imidazole. Purified PQBP1 constructs were then dialyzed into 20 mM Tris-HCl (pH 8.0) and 75 mM NaCl for 1 hr and were immediately used for TCCD measurements.

#### In vitro CA lattice assembly

Recombinant HIV-1 CA with and without the additional mutations (R18G, N74D) and CA K158C- AF568 were purified and labeled as described previously (Lau et al., 2019). CA lattices were assembled *in vitro* at 80 µM CA (1:99 ratio of CAK158C-AF568:CAA204C) in Tris-HCl (pH 8.0) and 1 M NaCl. The assembly solution was incubated 37°C for 15 min followed by 4°C overnight. Assembled CA lattices were dialyzed into 20 mM Tris-HCl (pH 8.0) and containing 75 mM NaCl.

#### Two-color coincidence detection (TCCD) spectroscopy

The TCCD fluorescence fluctuation spectroscopy approach has been described previously (Lau et al., 2021). Binding reactions were performed in 20 mM Tris-HCl (pH 8.0) and 75 mM NaCl containing 8 µM of CA (monomeric equivalent) that were assembled under high salt with binders as follows: 100 nM GFP-tagged PQBP1, 20 nM PQBP1-AF488, 50 nM CypA-AF488, 10 nM CPSF6_313-327_-AF488, and 10 nM of fluorescein-12-dATP (PerkinElmer, NEL465001EA). Where applicable hexacarboxybenzene (HCB) was added to a final concentration of 10 µM. At least 100 s of data were collected for each condition in 10 s acquisitions at 1000 Hz binning on an inverted microscope equipped with 488 nm (2.6 mW), 561 nm (0.5 mW) lasers and a water immersion 40x/1.2 NA objective (Zeiss). The emitted fluorescence from each fluorophore was separated into two channels using a dichroic mirror (565 nm) and filtered through a 525/50 nm band pass filter (AF488/GFP signal) and 590 nm long pass filter (AF568 signal), respectively, prior to focusing onto separate single photon avalanche diodes (Micro Photon Devices). The coincidence intensity ratios were calculated using a custom in-house software (TRISTAN, https://github.com/lilbutsa/Tristan/tree/master/v0_2/Matlab_Programs) as the slope of a curve obtained using weighted linear correlation of a graph plotting the binder fluorescence intensity plotting against its corresponding capsid signal intensities from each 10 s acquisition.(Lau et al., 2021) Coincidence ratios in different conditions were compared using ordinary one-way ANOVA using GraphPad Prism (v8.4).

**Video S1.**

**Time-lapse imaging to monitor structural integrity of an incoming viral particle with iGFP fluid phase marker (iGFP+IN-mRuby3) in MDDCs. See Figure S3B for detail.**

## SUPPLEMENTAL TEXT

### PQBP1 and IN distance distribution analysis

To assess the association, we analyzed the distances, *d*, from each IN to the nearest PQBP1 spot. The probability distribution of nearest neighbor distances, *P(d)*, shows a strong peak around *d*=0.125 µm, and a broader and weaker peak around *d*=0.7 µm (blue; left, Figure S1D). These two peaks likely represent two different aspects of protein localization. To distinguish between stochastic protein localization and biologically relevant colocalization, we manipulated the positions of IN and PQBP1 *in silico*. First, we randomized the labels, shuffling IN and PQBP1 spots together while keeping the density and IN/PQBP1 ratio constant. After label randomization, *P(d)* presents a single broad peak around *d*=0.75 µm (purple; left, Figure S1D). Next, we added a random uniform jitter *J* to PQBP1 positions and calculated *P(d)*. We find that any added jitter weakens the narrow peak at *d*=0.125 µm in favor of the peak at *d*=0.75 µm. At *J*=1 µm the jittered curve (yellow; left, Figure S1D) overlays of the randomized curve. One way to quantify colocalization is to calculate the percentage *ϕ* of IN spots that have a PQBP1 within a chosen threshold, *δ* (right, Figure S1D). Formally, *ϕ* is defined by 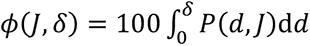. A reasonable value for *δ* accounts for the optical aberrations, which can be wavelength dependent. While the two fluorophores may be physically very close, their image on a microscope could be shifted and deformed. The safest way to proceed is then to choose a reasonable value for *δ* and calculate the colocalization using that threshold. Finally, the resistance to one’s findings to small variations in *δ*, known as sensitivity analysis, can be used to support the original choice of parameter. We plot *ϕ* as a function of jitter *J* and note that for the reasonable values of *δ* (0.1 µm & *δ* & 0.75 µm ), the dependence of *ϕ* on *J* follows a similar behavior, implying that δ is robust against small changes. The biphasic decrease in *ϕ* again indicates the presence of two aspects of protein colocalization. We note that the step in *ϕ* happens when *J∼δ*. This does not occur when the labels are randomly shuffled, indicating that once the jitter exceeds the nearest neighbor distance, the colocalization is destroyed in real data, but not in randomized shuffling on acquired data. Collectively, the distance analysis confirmed a robust co-localization between PQBP1 protein and incoming virions (above 40% at *δ*, = 0.5 µm) that is not measured due to random distribution of cellular PQBP1.

## SUPPLEMENTAL LEGENDS

**Figure S1.**
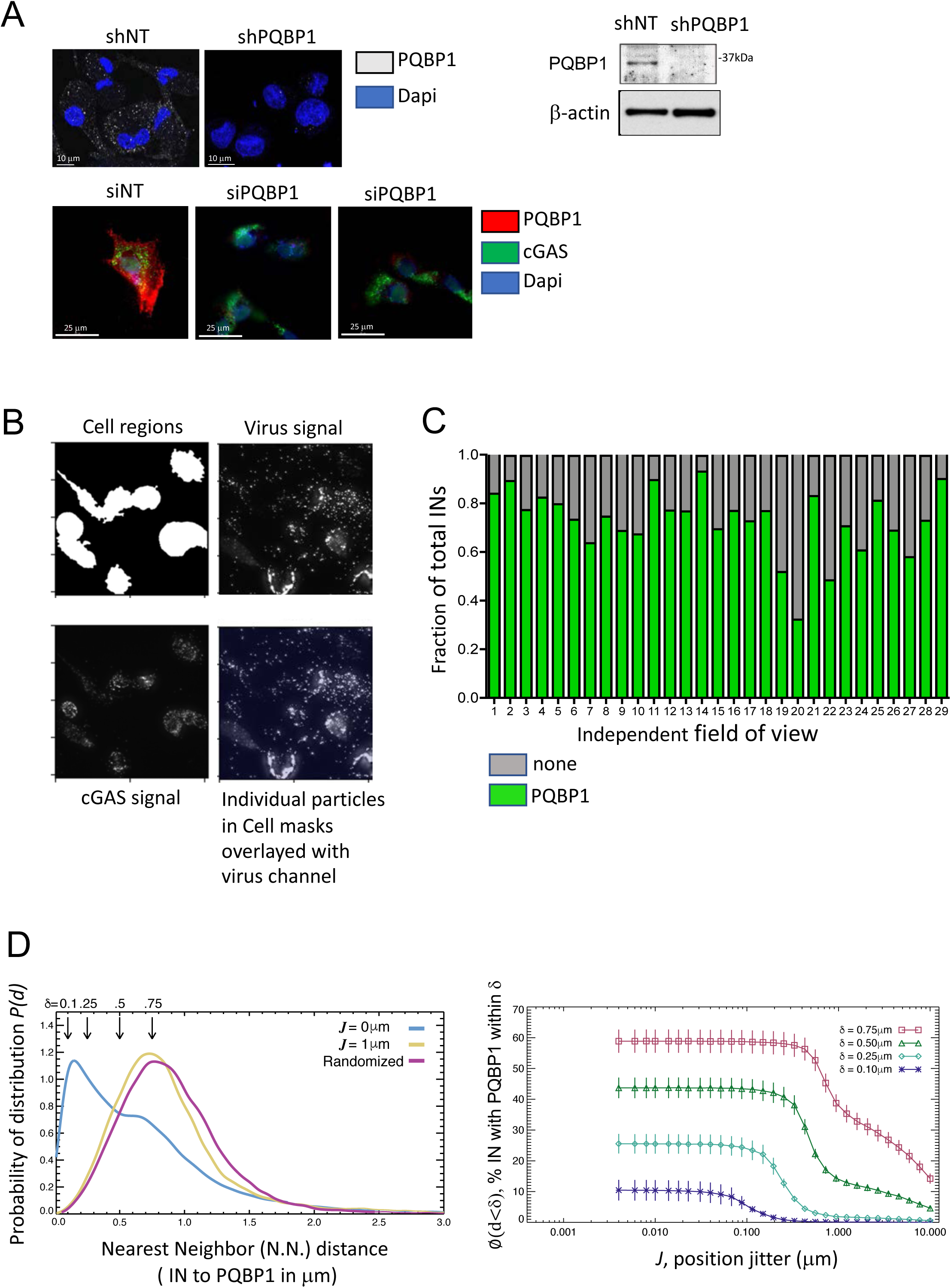
PQBP1 association with incoming HIV-1 virus particles. (A) Specificity of PQBP1 antibody (Sigma) and lack of effect on overall cGAS expression was confirmed by IF and/or western blot of THP-1 cells expressing indicated shRNAs (top) or MDDCs treated with indicated siRNAs (bottom). (B) Identification of viral particles from infected THP-1 cells was done with python with skimage, scipy, pims, and trackpy tools. In short, cells were subjected to a binary threshold so that no extracellular signals (viruses or cell debris) would confound the analysis. Viral particles were detected using thresholding and watershed segmentation methods or by using trackpy. The masks of the pixels that are positive for viral particles are then quantified for each independent channel (PQBP1, cGAS, iGFP, IN label, etc) and (C) for each field of view that was imaged, fraction of total virions (IN) showing positive PQBP1 signal (compared to secondary antibodies-only controls) were graphed. (D) Quantification of PQBP1 and IN distance distribution analysis. Left, probability distribution of distances, *P(d)*, from each virion (IN) to the nearest PQBP1 (blue). *P(d)* for uniformly distributed jitter *J=1 µm* (yellow) becomes similar to the distribution obtained by random shuffling of puncta (purple). Arrows indicate reasonable arbitrary thresholds δ to be applied to *d* values in determining association of PQBP1 puncta with corresponding IN. Data were obtained from 24 independent images (N=24,471). Right, the percentage of IN spots that have a PQBP1 within a chosen threshold, *δ* (0.1 µm < *δ* < 0.75 µm). See Supplemental Text for detail.

**Figure S2.**
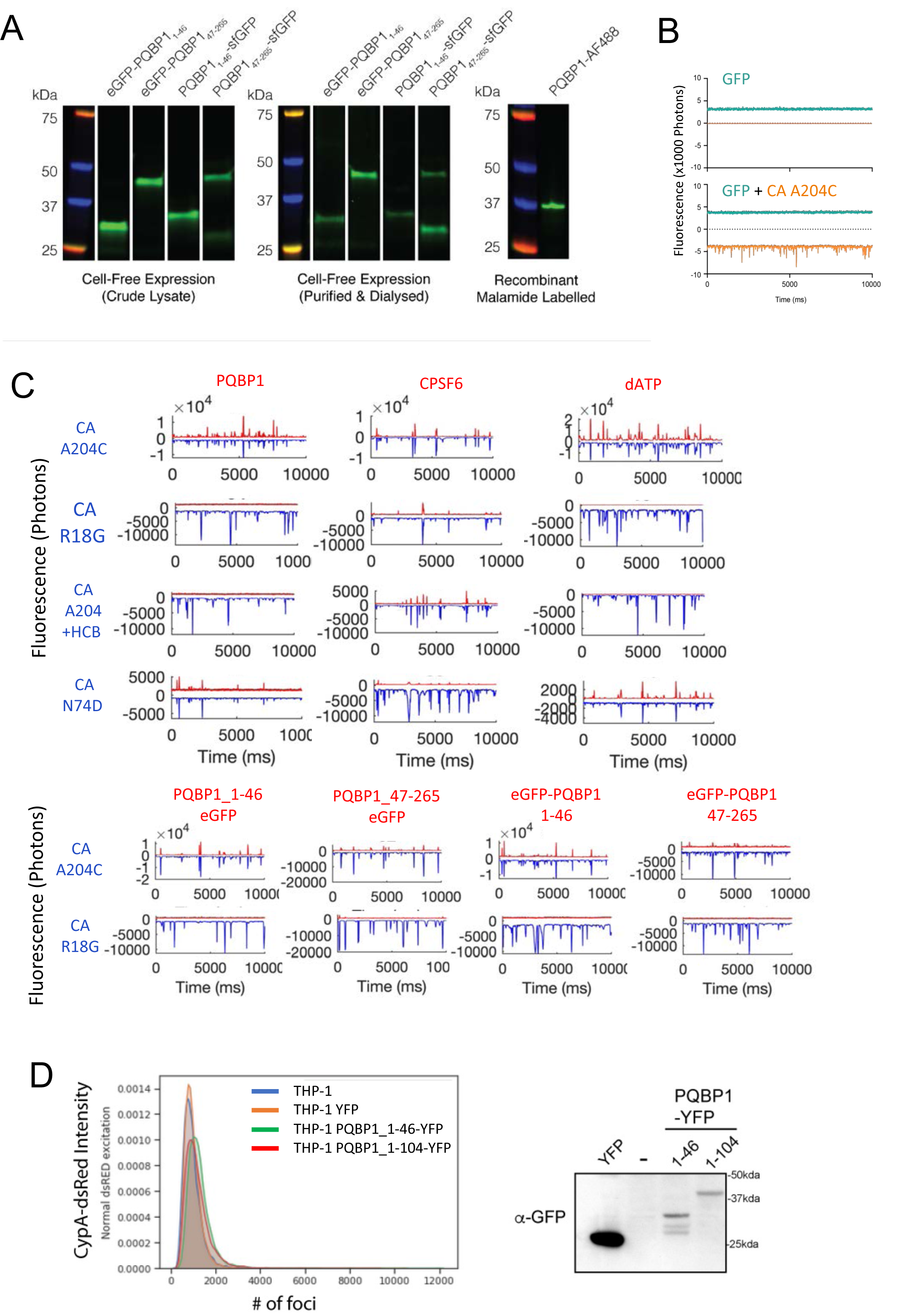
Mapping of capsid interaction domain in PQBP1. (A) PQBP1_1-46_ and PQBP1_47-265_ with N-terminal eGFP or C-terminal sfGFP produced in a cell-free protein expression mixture before (left) and after (middle) purification by Ni-NTA chromatography followed by dialysis. Recombinant PQBP1 was labelled at C60 with AF488-maleimide (right). (B) Dual-color fluorescence traces of a negative control GFP (green traces) with AF568-labelled CA A204C particles (orange traces). (C) Top panels, representative fluorescence traces of either AF488 labelled PQBP1, CPSF6 peptide or dATP (red traces) and the CA A204C particles (blue traces). Inclusion of hexacarboxybenzene (HCB) or use of either N74D or R18G double mutants with CA A204C is indicated. Bottom panels, analogous to the top, except eGFP fused PQBP1 truncation constructs were utilized. (D) Fluorescence resonance energy transfer (FRET) assay to measure interaction between PQBP1_1-46_-YFP and CypA-dsRed. THP-1 cells either wild type or stably expressing either eYFP, PQBP1_1-46_-eYFP or PQBP1_1-104_-eYFP were infected with HIV-1 virus packaged with CypA-DsRed for 1.5 hrs, followed by PFA fixation and FRET analysis. Signal intensity distribution of CypA-dsRed positive foci for indicated THP-1 clones infected with the virus (left) and a western blot depicting expression of both eYFP and PQBP1-eYFP proteins are shown.

**Figure S3.**
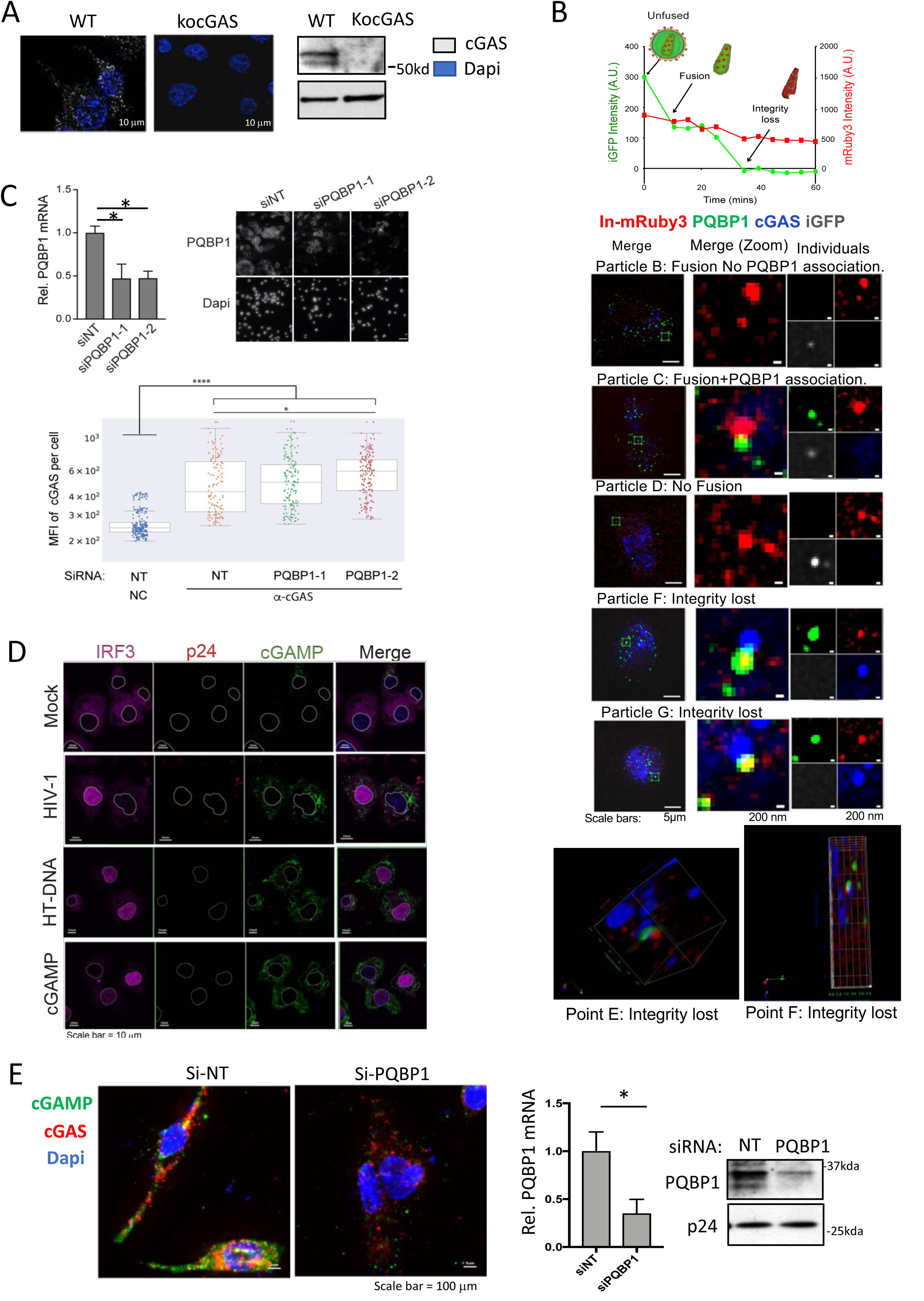
PQBP1 is required for cGAS recruitment to incoming virus particles. (A) The specificity of cGAS antibody was confirmed by IF and western blot on cGAS knock out in THP-1 cells. (B) Time lapsed imaging to monitor the structural integrity of an incoming viral core. MDDCs are infected with HIV-1 virus with iGFP fluid phase marker (iGFP+IN-mRuby3 at low MOI) where iGFP and IN-mRuby signal intensities of each virion were monitored for one hour. See Video S1. A representative kinetic profile of the signal intensities of a virion at different stages where the loss of GFP signal linked to virion fusion and capsid integrity loss are depicted (top). Panels of each virion, identified by both time-lapsed and corresponding IF x-y-z position of the respective foci point label was categorized based on the capsid integrity status and are color coded for the signals of PQBP1 (green), cGAS (blue), IN-mRuby3 (red). The RGB merged images and iGFP (gray in the individual boxes) are also shown (middle, particle B to G). 3D reconstruction of point E and point F foci are shown where we see the proximity and overlapping of PQBP1/cGAS complexes to the viral particle. (C) A knock down efficiency of siRNAs against PQBP1 is demonstrated by both RT-qPCR and IF analysis (left and right respectively). MFI of cGAS signal intensity per THP-1 cell treated with either non-targeting (NT) or PQBP1 specific siRNAs and infected with HIV-1 virus as described in Figure 3C. Mean and error bar (-/+ 1.5*IQR) are shown. Boxes denote the interquartile range. T-test (two-tailed, equal variation), *p<0.05; Dunn’s column comparison **** p<0.0001, * p<0.05. (D) Increase in cGAMPs in the cytosol of the infected cells. PMA-THP-1 cells, either mock infected or infected with HIV-1 luciferase virus or transfected with either HT-DNA or cGAMP for 3 hours, were stained for cGAMP (green), p24 (red), IRF3 (magenta) and dapi (blue). The white contour line indicates the boundary of nucleus. (E) Knockdown of PQBP1 results decreased cGAMP production. Left, PMA-THP-1, transfected with non-targeting siRNAs (NT) or siRNA targeting PQBP1, were challenged with HIV-1, stained for cGAMP (green), cGAS (red), and dapi (blue). Right, knockdown efficiencies of both mRNA and protein level of PQBP.

**Figure S4.**
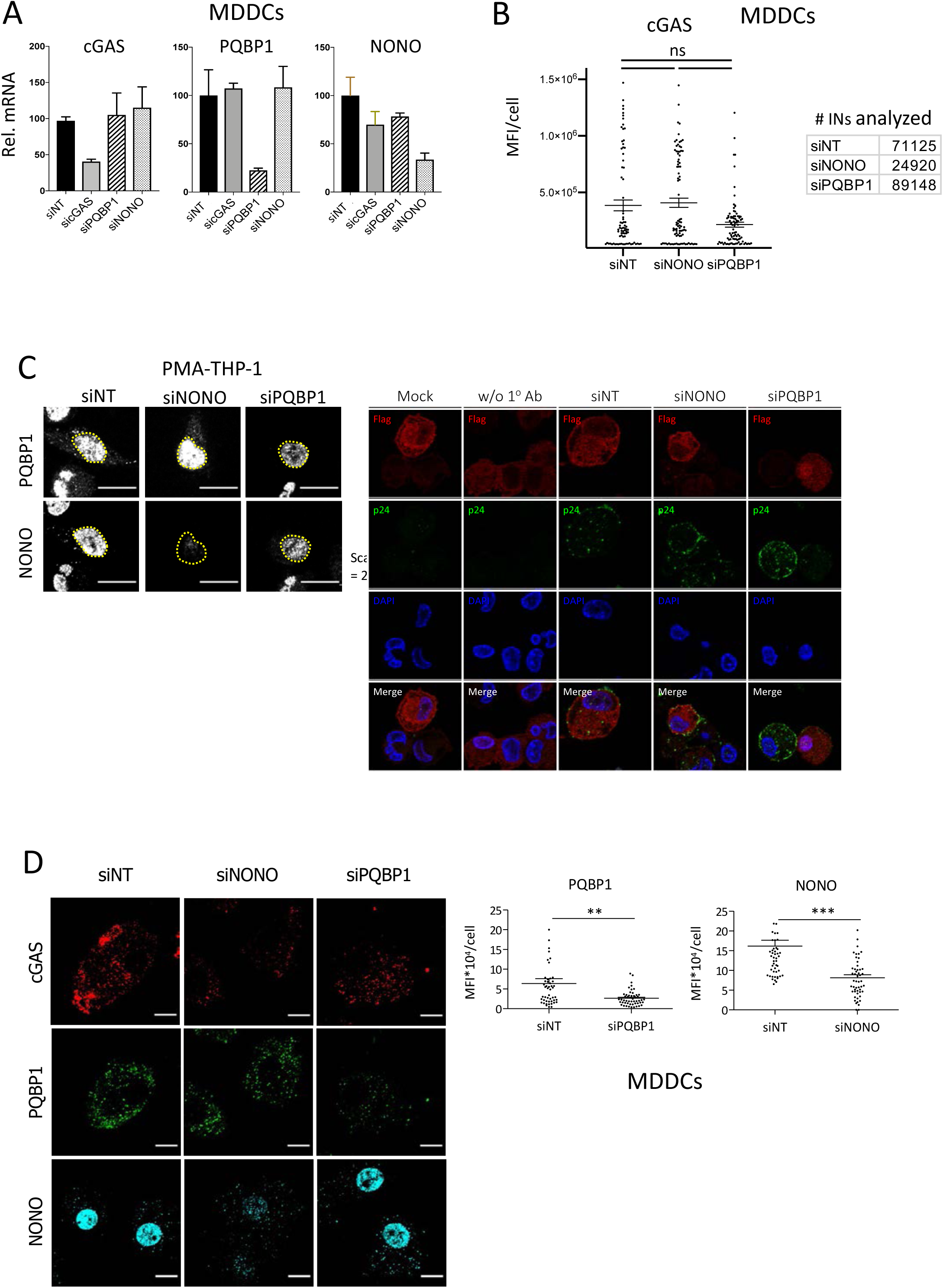
NONO is not required for the PQBP1-dependent cGAS sensing occurring during the early step of the infection. (A) PMA differentiated THP-1 cells, subjected to siRNA-mediated targeting were infected with were with either mock of HIV-1 in the presence of VLP-Vpx for 2 hrs, followed by post-fixation IF imaging. The level of NONO and PQBP1 proteins of the indicated THP-1 cells (top) and expression levels of Flag-cGAS (red), p24 (green) and Dapi (blue) of the cells utilized for proximal ligation assay as in Figure 4 (bottom) are shown. (B) Knockdown efficiencies of siRNA-targeted genes in MDDCs were quantified by RT-qPCR. (C) MFIs of cGAS signal per infected MDDC, subjected to indicated siRNA treatments, are shown. A table at the bottom of the graph shows # of INs analyzed for each condition. (D) Representative images of the infected MDDCs having treated with indicated siRNAs (left) and MFI of indicated protein signals per cells (right) analyzed in Figure 4D and 4F are shown. Mean and SEMs are shown. One-way ANOVA, ***p<0.001, **p<0.01, *p<0.05. ns denotes no significance.

**Figure S5.**
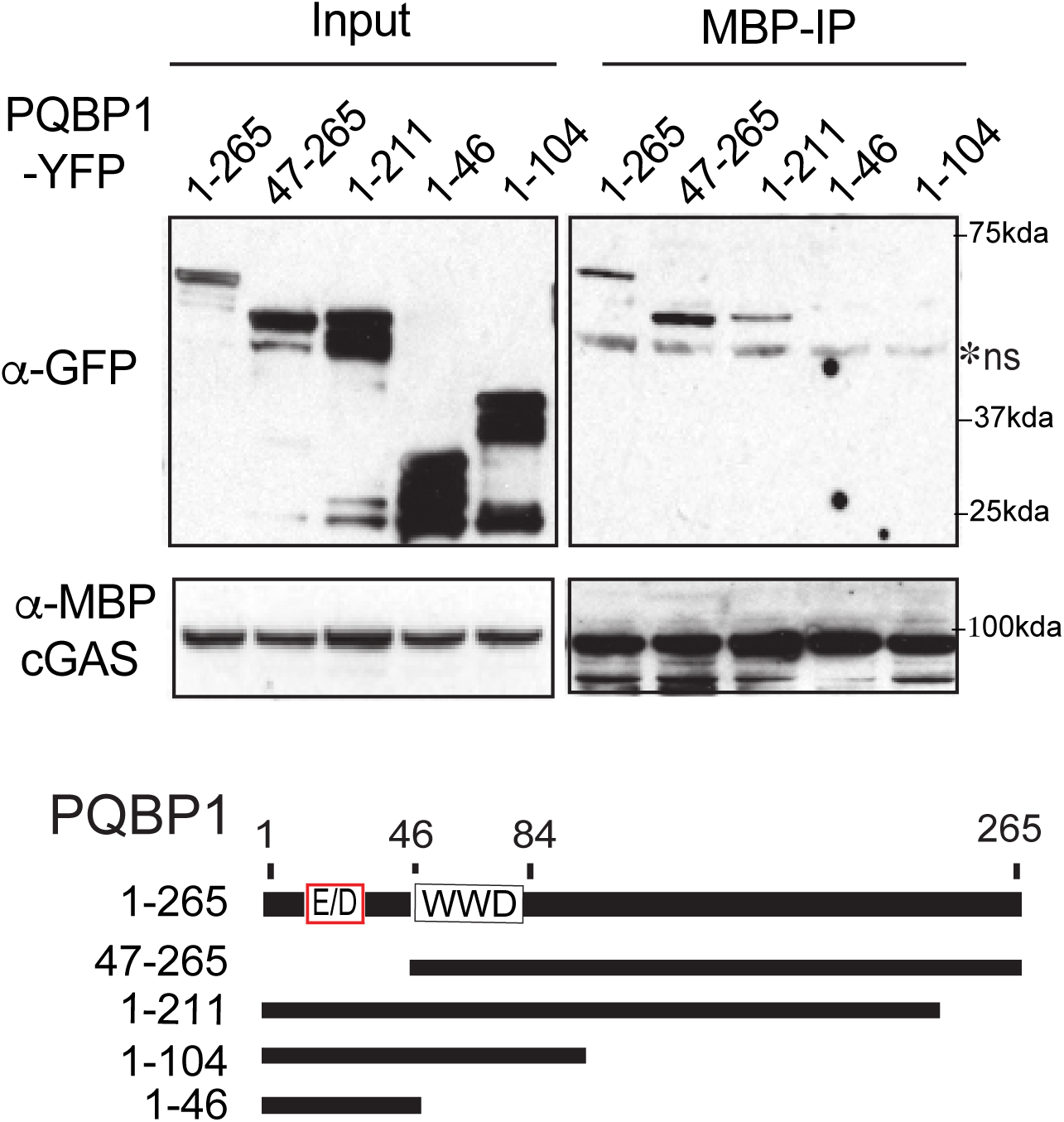
N-terminal capsid interaction domain of PQBP1 is dispensable for cGAS interaction. Either full-length or truncated PQBP1-YFP proteins were co-expressed with MBP- cGAS in 293T cells and subjected to anti-MBP pull down. *ns denotes non-specific protein. A schematic of PQBP1 protein is shown. WWD and E/D denote ww domain and acidic aa rich domain respectively.

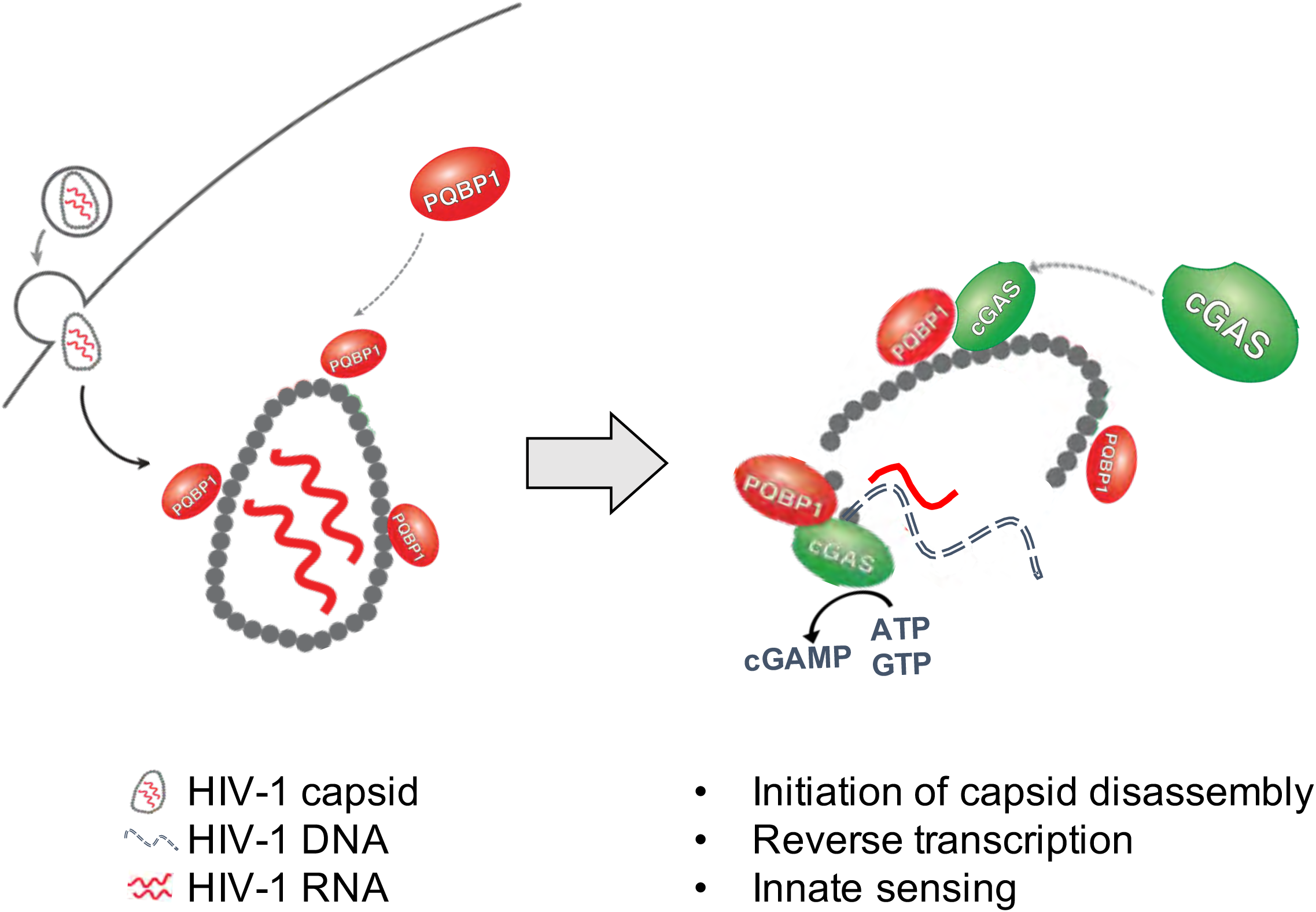

